# AT6SS trans-kingdom effector is required for the delivery of a novel antibacterial toxin in *Pseudomonas aeruginosa*

**DOI:** 10.1101/551929

**Authors:** Benjamin Berni, Chantal Soscia, Djermoun Sarah, Ize Bérengère, Sophie Bleves

## Abstract

*Pseudomonas aeruginosa* has evolved multiple strategies to disarm and take advantage of its host. For this purpose this opportunist pathogen has particularly developed protein secretion in the surrounding medium or injection into host cells. Among this, the Type VI Secretion System (T6SS) is utilized to deliver effectors into eukaryotic host as well as target bacteria. It assembles into a contractile bacteriophage tail-like structure that functions like a crossbow, injecting an arrow loaded with effectors into the target cell. The repertoire of T6SS antibacterial effectors of *P. aeruginosa* is remarkably broad to promote environmental adaptation and survival in various bacterial communities, and presumably in the eukaryotic host too.

Here we report the discovery a novel pair of antibacterial effector and immunity of *P. aeruginosa*, Tle3 and Tli3. Tli3 neutralizes the toxicity of Tle3 in the periplasm to protect from fratricide intoxication. The characterization of the secretion mechanism of Tle3 indicates that it requires a cytoplasmic adaptor, Tla3, to be targeted and loaded onto the VgrG2b spike and thus delivered by the H2-T6SS machinery. Tla3 is different from the other adaptors discovered so far, and defines a novel family among T6SS.

Interestingly this led us to discover that VgrG2b that we previously characterized as an anti-eukaryotic effector possesses an antibacterial activity as well, as it is toxic towards *Escherichia coli*. VgrG2b is thus a novel trans-kingdom effector targeting both bacteria and eukaryotes. VgrG2b represents an interesting target for fighting against *P. aeruginosa* in the environment and in the context of host infection.

**Highlights:**

- Tle3 and Tli3 are a novel pair of antibacterial toxin and immunity of *P. aeruginosa*
- Tla3 recruits Tle3 in the cytoplasm, and targets it to VgrG2b
- VgrG2b is required for Tle3 delivery into target bacteria by the H2-T6SS
- Tla3 defines a novel type of T6SS adaptor with a DUF2875
- VgrG2b is a new trans-kingdom effector targeting both bacteria and eukaryotes

## Introduction

*Pseudomonas aeruginosa* is one of the most virulent opportunistic pathogens, being responsible for various diseases such as acute infections of lungs and burned skin that can lead to septicemia more particularly in immunocompromized patients, or broncho-alveolar colonization in Cystic Fibrosis sufferers. *P. aeruginosa* has been classified in 2017 as critical by the WHO in the top three list of antibiotic resistant bacteria [1]. *P. aeruginosa* has developed various pathogenicity strategies among which protein secretion or protein delivery into target cells is key. Indeed this pathogen possesses 5 of the 6 secretion systems so far identified among Gram-negative bacteria, if we exclude the T9SS (Type IX Secretion System) restricted to one phylum, and remarkably in several copies for most of them [2].

The T6SS (Type VI Secretion System) was first discovered in the context of eukaryotic host infection [3][4] and later during bacterial competition [5], which seems to be its primary function [6]. The T6SS confers a fitness advantage (i) in environmental niches against rival bacteria (inter- and intraspecies competitiveness have been described) and (ii) in the eukaryotic host towards commensal bacteria [6]. Indeed recent studies have highlighted a novel role for T6SS-dependent antibacterial responses in interbacterial competition in the mammalian gut [7][8], suggesting that T6SSs may be important not only in shaping microbial community composition, but also in governing interactions between the microbiota and invading pathogens. Interestingly, several T6SSs are also known to target both cell type genus such as the T6SS of *Vibrio cholera* [4][9] and *P. aeruginosa* [10][11]. Even more remarkably, three T6SS effectors of *P. aeruginosa*, namely PldA (also called Tle5a), PldB (Tle5b) and TplE (Tle4) [12,13] have been called “trans-kingdom effectors” since these toxins can target both prokaryotic and eukaryotic cells [14]. Indeed toxins are usually directed against eukaryotic cells (like AB toxins or RTX pore-forming toxins) or against rival bacteria (like bacteriocins).

The T6SS functions as a dynamic contractile phage tail-like structure anchored in the bacterial cell envelope that delivers effector proteins directly into the target cell in a one-step manner. T6SS includes a contractile sheath that cover a nanotube of stacked Hcp topped with a membrane-puncturing spike made of VgrG and PAAR (proline-alanine-alanine-arginine repeat) proteins [15]. The sheath can contract and inject the arrow loaded with effectors into the target cell. Characterizing the repertoire of effectors delivered by the T6SS has highlighted a great diversity in terms of effector activities, host cell targets, and mode of recruitment by the T6SS machinery. In brief, there are two broad effector categories: the “specialized” effectors fused to components of the machinery (evolved VgrG, evolved PAAR and evolved Hcp have been described so far) and the “cargo” effectors [6]. The later are addressed to the T6SS machinery by binding directly one of the arrow components (VgrG, PAAR and Hcp) or by being targeted through cytoplasmic adaptor proteins also called chaperones. To date three families of adaptor have been described, the first one harboring a DUF4123 [16–19], the second a DUF1795 [20,21] and the third one a DUF2169 [22]. In line with this, many effector-encoding genes are found in close proximity to *vgrG, hcp, paar* or adaptor genes. Finally to protect themselves from self-intoxication or from antibacterial toxins injected by neighboring sibling cells, bacteria always synthesize immunity proteins, which are encoded by adjacent genes [23].

*P. aeruginosa* encodes three distinct T6SS loci, H1- to H3-T6SS. While H1-T6SS is only involved in antibacterial activity so far [24][25], H2-T6SS and H3-T6SS can target both bacterial and eukaryotic cells possessing even as said earlier trans-kingdom effectors [10–13,26,27]. We discovered the anti-eukaryotic function of the H2-T6SS machinery that promotes the uptake of *P. aeruginosa* by non-phagocytic cells [10]. The two phospholipases D mentioned earlier, PldA (Tle5a) and PldB (Tle5b), delivered respectively by H2-T6SS and H3-T6SS machineries, participate in the host kinase pathway highjacking that facilitates further entry of *P. aeruginosa* [12]. The evolved VgrG2b effector [26] is delivered by H2-T6SS into epithelial cells where it targets the γ-tubulin ring complex, a microtubule-nucleating multiprotein complex to promote a microtubule-dependent internalization of *P. aeruginosa*. Finally TplE (Tle4), which is secreted by the H2-T6SS machinery, promotes autophagy in epithelial cells once localized to the endoplasmic reticulum [13]. Interestingly PldA (Tle5a), PldB (Tle5b) and TplE (Tle4) have also been identified as antibacterial phospholipases of the Tle (type VI lipase effectors) family [11]. They work by affecting membrane integrity of the rival bacteria [11–13]. More precisely, PldA degrades the major constituent of bacterial membranes, the phosphatidylethanolamine [11].

In the present study we have discovered a novel antibacterial toxin, Tle3 and its cognate immunity, Tli3, whose genes are encoded downstream of *vrgG2b*. By characterizing the secretion mechanism of Tle3 by H2-T6SS, we showed that it requires Tla3, a cytoplasmic adaptor of a unique family, to be targeted to the VgrG2b spike. Interestingly we also found that the C-terminal extension of VgrG2b is toxic towards *Escherichia coli* making VgrG2b a new trans-kingdom effector of *P. aeruginosa*.

## Results

### Tle3 is a novel antibacterial toxin of ***P. aeruginosa***

The analysis of the *vgrG2b* genetic environment revealed the presence of the PA0260 gene encoding a protein with a α/β hydrolase domain (PF00561) and a putative Ser-Asp-His catalytic triad used by various esterase enzymes and has thus been classified in the Tle3 family of antibacterial Tle toxins [11] (Fig. 1A). The immunities of Tle proteins, which are lipolytic toxins active in the periplasm of the prey bacterium, are localized in or exposed to the periplasm where they neutralize the cognate toxin [11–13,29]. The two genes surrounding *tle3*, PA0259 and PA0261, are good candidates as Tle3 immunity. Indeed sequence comparisons of PA0259 and PA0261 showed that PA0261 is homologous to Tsi6, the immunity protein of a H1-T6SS effector called Tse6, and PA0259 to TplEi (Tli4), the immunity of TlpE (Tle4), a H2-T6SS effector [13,30]. We used the SignalP 4.1 server [31] to predict the cellular localization of the two immunity candidates. While a Sec signal sequence is predicted at the N-terminal extremity of PA0261, the analysis of the PA0259 sequence did not reveal any. However three upstream ATG can be found in frame with the annotated ATG of PA0259 (Fig. 1 sup). The sequence of the proteins synthesized from the two first ATG (ATG_1_ et ATG_2_) presents then a N-terminal signal peptide, while the protein synthesized from the last codon (ATG_3_) does not. Moreover a RBS (Ribosome Binding Sequence) can be found only upstream of ATG_1_ and with a significant Kolaskar score that indicates a strong probability to be used as an initiation codon [32]. Altogether these data tend to indicate an incorrect annotation of the start codon of PA0259 and that ATG_1_ should be considered for the initiation of PA0259 translation. Such a correction has been already seen for Tli5 (PA3488) the immunity protein of PldA (Tle5a) of *P. aeruginosa* [11]. In conclusion of this *in silico* analysis, the two putative immunities may harbor a Sec signal peptide that suggests a periplasmic localization in agreement with Tle3 activity in this compartment.

**Fig. 1:**
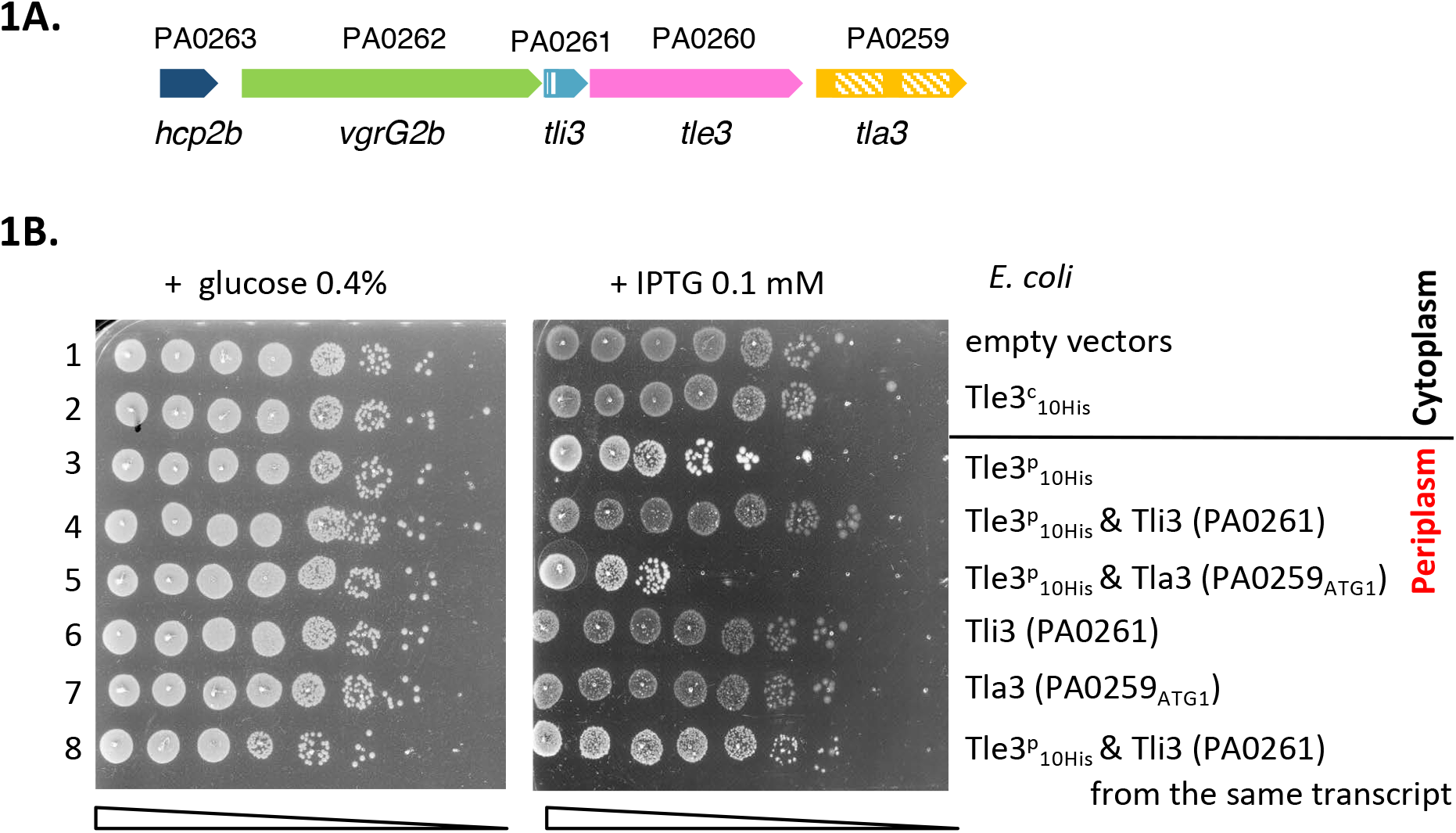
***vgrG2b* island organization (A)** The genes are labeled with the given name (i.e., *hcp2b*) and are indicated by their annotation number (e.g., PA0263). The Sec signal peptide of Tli3 (PA0261) and the two DU2875 of Tla3 (PA0259) are represented with stripped boxes. **The Tle3 periplasmic toxicity is counteracted by Tli3 (PA0261) (B).** Serial dilutions (from non-diluted to 10^−7^) of normalized cultures of *E. coli* BL21(DE3)pLysS producing the wild-type Tle3 in the cytoplasm, called Tle3^c^ (from pVT1, a pETDuet-1 derivate) or in the periplasm, called Tle3^p^ (from pSBC81, a pET22b(+) derivate yielding a fusion of Tle3 with a Sec signal peptide) were spotted on LB agar plates supplemented (left panel) with 0.4% glucose or (right panel) with 0.1mM IPTG. Glucose and IPTG allow respectively repression and induction of the gene encoding the T7 RNA polymerase. When indicated Tli1 (PA0261) or Tla3 (PA0259) were produced in the periplasm from pVT8, pVT9 and pSBC107 respectively, pRSFDuet-1 derivates. Line 1: pET22b(+) and pRSFDuet-1, line 2: pVT1 and pRSFDuet-1, line 3: pSBC81 and pRSFDuet-1, line 4: pSBC81 and pVT8, line 5: pSBC81 and pVT9, line 6: pVT8 and pET22b(+), line 7: pVT9 and pET22b(+) line 8: pSBC107 is a pRSFDuet-1 derivate producing Tli3 (PA0261) and Tle3^p^ from the same transcript.

To demonstrate the antibacterial activity of Tle3 and to identify its immunity protein, we developed a heterologous toxicity assay in *E. coli* on the basis that Tle3 should be toxic when produced in the periplasm of *E. coli* and that it should be counteracted by the co-production of its immunity protein. In order to artificially address Tle3 to the periplasm of *E. coli* the *tle3* sequence has been cloned in frame with the sequence coding the PelB signal peptide on the pET22b vector under a P_T7_ promoter. PA0259 from ATG_1_ and PA0261 were respectively cloned in pRSF-DUET, a higher copy-number vector, to allow a maximal co-expression with *ss-tle3* in the *E. coli* BL21(DE3) pLysS strain. The correct production and localization of all the recombinant proteins in *E. coli* has been verified by western blot after cell fractionation (Fig. 2 sup). The results presented in Figure 1B indicate that whereas the cytoplasmic production of Tle3 was not toxic, Tle3 targeting to the periplasm led to *E. coli* killing. Moreover while PA0259 had no effect, the coproduction of PA0261 in the periplasm protected the cells against the toxicity of Tle3. We verified that the sole overproduction of PA0259 and PA0261 was not toxic to *E. coli*. As PA0261 neutralized Tle3 toxicity, we called it “Tli3” for Type VI lipase immunity (Fig. 1A). During this study, we observed that the protection conferred by Tli3 coproduction could be sometimes partial and we solved this issue by cloning in tandem *tli3* and *tle3* on the same plasmid like they are organized on *P. aeruginosa* genome (Fig. 1B, line 8). The importance of this genetic link is the proof of the close connection between these two proteins as a pair of toxin-antitoxin.

**Fig. 2:**
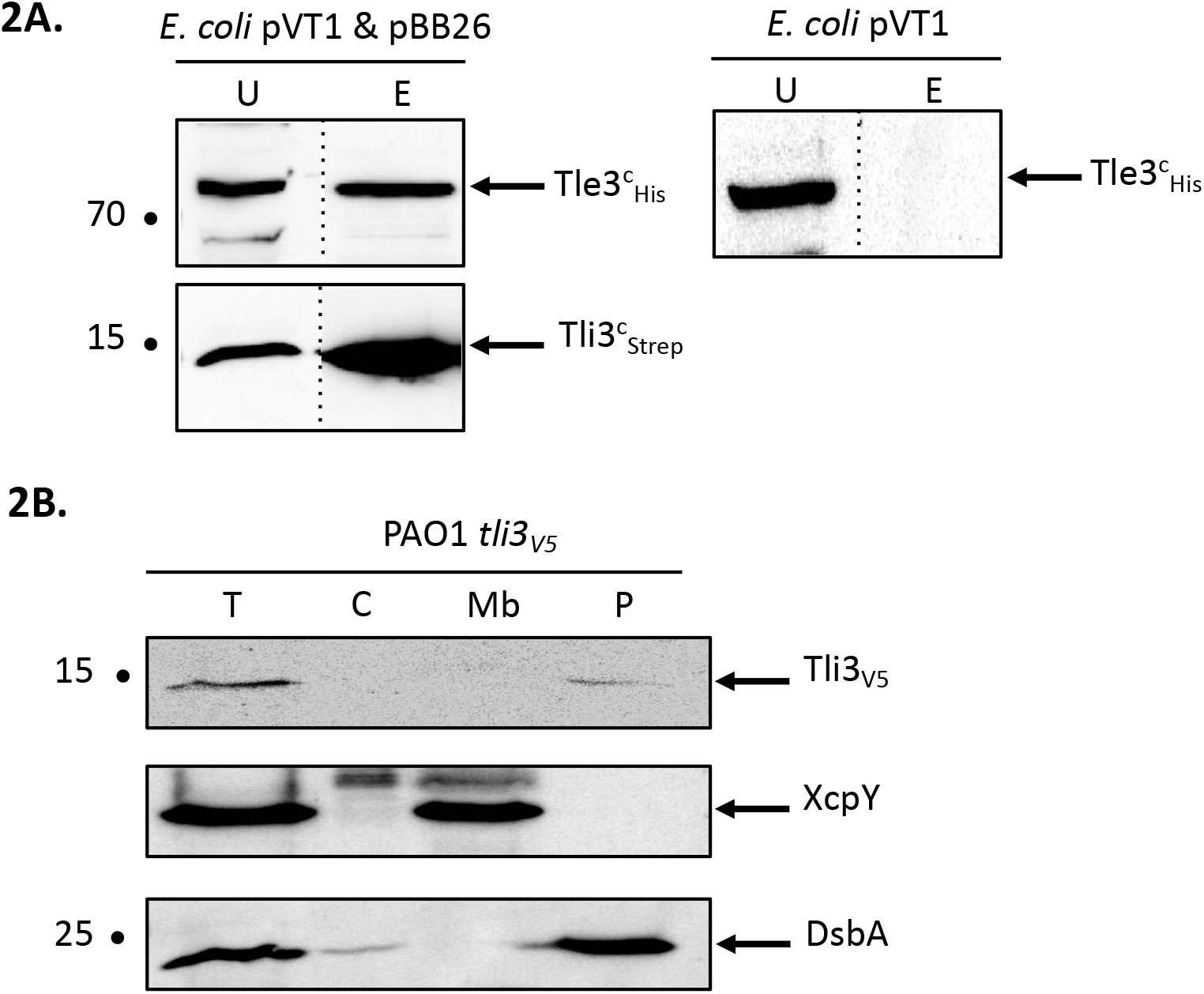
**Tli3 (PA0261) interacts with Tle3 (A)** Copurification assay on StrepTactin column of Tli3cSTREP with Tle3c10HIS produced in *E. coli* BL21(DE3)pLysS produced from pBB26 and pVT1 respectively. The unbound (U) and eluted (E) fractions were collected and subjected to SDS-PAGE (10.5 %) and Western blot analyses using anti-His antibody (Upper) and anti-streptavidin antibody (Lower). The position of the proteins and the molecular mass markers (in kDa) are indicated. **Tli3 (PA0261) is a periplasmic protein in *P. aeruginosa* (B)** Cells of *P. aeruginosa* PAO1 tli3V5 were subjected to fractionation and immunoblotting using antibodies directed against the V5 tag, XcpY and DsbA. XcpY and DsbA were used as membrane and periplasmic controls respectively. T: whole cell, C: cytoplasm, Mb: total membrane, P: periplasm. The position of the proteins and the molecular mass markers (in kDa) are indicated.

### Tli3 (PA0261) is the immunity protein of Tle3

As the immunities bind specifically their effector, which is suggested by the release of Tle3 toxicity by Tli3, we tested the physical interaction between both proteins by co-purification with affinity chromatography (Fig. 2A). A cytoplasmic Strep-tagged version of Tli3 was engineered by fusing the tag to the mature domain of Tli3, lacking its signal peptide. The recombinant protein was coproduced in *E. coli* BL21(DE3) pLysS with the cytoplasmic His_10_-tagged Tle3. The bacterial lysate was loaded to a StrepTactin matrix (see Materials and Methods section), and Tli3^C^_Strep_ was eluted with desthibiotin. The presence of Tle3^C^_His_ was controlled in the elution fraction with anti-His antibodies. As showed in Fig. 2A, Tle3^C^_His_ was found in the eluted fraction only upon coproduction with Tli3^C^_Strep_ (left panel). Indeed when produced alone in *E. coli* Tle3^C^ _His_ was not purified by affinity chromatography (right panel). As expected for an immunity protein, Tli3 directly interacts with Tle3.

To go further into Tli3 characterization we chose to determine its cellular localization in *P. aeruginosa*. All the immunity proteins identified so far for Tle proteins are localized in the periplasm or associated to the periplasmic side of the outer membrane [11–13,29] in order to counteract their cognate toxin. A chromosomally encoded Tli3_V5_ translational fusion was engineered in order to specifically immunodetect the protein in *P. aeruginosa* (Supplementary Table 1). After fractionation of *P. aeruginosa* (Fig. 2B), Tli3 was readily observed in the same fraction as DsbA that catalyzes intrachain disulfide bond formation as peptides emerge into the periplasm. This indicates a periplasmic localization for Tli3 in *P. aeruginosa* in agreement with the presence of a Sec signal peptide and our working hypothesis suggesting an immunity function.

### Tle3 interaction network

To further characterize Tle3, we performed a bacterial two-hybrid (BACTH) assay with the other gene products of the *vgrG2b* operon hypothesizing that a genetic link could reflect protein-protein interaction. The sequences coding PA0259 and Tli3 (PA0261) after their signal sequences and Tle3 were cloned downstream and upstream the T18 or T25 domains of the *Bordetella adenylate* cyclase. Because of the high molecular weight of VgrG2b and since the interaction of another Tle with a VgrG in entero-aggregative *E. coli* (EAEC) was previously delimitated to the C-terminal domain of VgrG [29], we cloned the sequences encoding the C-terminal extension (domains 1, 2 and 3) and a truncated version harboring only the DUF2345 and the transthyretin-like (TTR) domains (domains 1 and 2) of VgrG2b downstream the T18 and T25 domains (Fig. 3A).

**Fig. 3:**
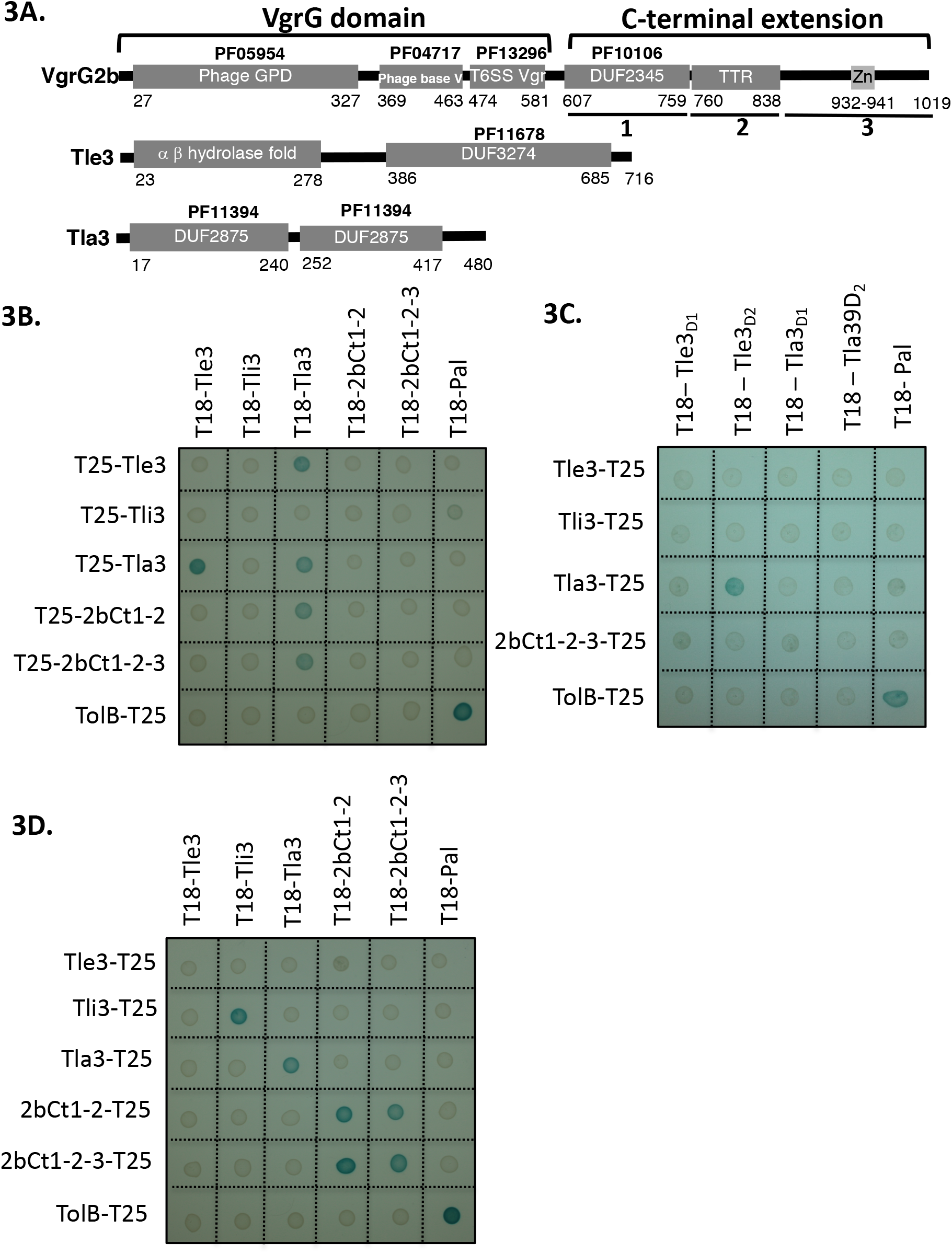
**Tla3 (PA0259) interacts with the Tle3 toxin and with VgrG2b. Domain organization of VgrG2b, Tle3 and Tla3 (PA0259) (A).** The first 581 residues of VgrG2b carry the VgrG domain homologous to gp27 and gp5 phage tail proteins and consisting of three sub-domains. This is followed by the C-terminal extension, composed of a conserved domain of uncharacterized proteins (DUF2345, PF10106) (domain 1), a TTR (transthyretin-like region) (domain 2), and a putative zinc-dependent metallopeptidase pattern (LFIHEMTHVW signature, PS00142 (in domain 3). Tle3 architecture consists of an α β hydrolase fold domain followed by a DUF3274, Tla3 of a tandem of DUF2875. **Bacterial two-hybrid assay (B, C and D).** BTH101 reporter cells producing the indicated proteins or domains fused to the T18 or T25 domain of the *Bordetella* adenylate cyclase were spotted on X-gal indicator plates. The blue color of the colony reflects the interaction between the two proteins. TolB and Pal are two proteins known to interact but unrelated to the T6SS. The experiment was performed in triplicate and a representative result is shown.

Unexpectedly the sole interaction revealed by the BACTH assay for Tle3 was with PA0259 (Fig. 3B) since only the T18/T25-Tle3 and T18/T25-PA0259 fusion proteins coproduction activated the expression of the reporter gene. This assay did not confirm the interaction between Tle3 and Tli3 observed by copurification, and did not evidence an interaction with VgrG2b. Interestingly Tle3 and PA0259 did not interact anymore when they were fused upstream the T18 and T25 domains suggesting that both proteins interact via their C-terminal domains (Fig. 3A Sup). To go further in characterizing the interaction between Tle3 and PA0259, we constructed truncated variants taking into account their domain organization (Fig. 3 A). We further delimitated the interacting domain of Tle3 to its extreme C-terminus (the DUF3274 domain) since only the truncated T18-Tle3D2 construct still interacts with T25-PA0259 (Fig. 3C). We did not find the domain of interaction within PA0259 as none of the DUF2875 domains alone interacts with Tle3 or this may suggest that both of them are required for the interaction (Fig. 3C).

Next, we took advantage of all the constructs we made to test other interactions. The BACTH assay also showed that PA0259 interacts with both forms of the VgrG2b C-terminal extension (Fig. 3B). The domain of interaction on VgrG2b is thus at least constituted by the DUF2345 and TTR domains. We also observed that PA0259 at least dimerizes since all the PA0259 constructs interact with each other whatever the orientation of PA0259 (Fig. 3B, 3D, Fig. 3B Sup). VgrG2b and Tli3 (PA0261) are also able to homomultimerize since all the constructs interact with each other (Fig. 3D, Fig. 3B Sup).

Taking into account the interactions revealed by the BACTH assay, we propose that the Tle3 toxin can be addressed to the H2-T6SS machinery VgrG2b component via PA0259. We thus named PA0259 “Tla3” for Type VI lipase adaptor protein.

### Tla3 (PA0259) characterization

To gain insight into the role of Tla3 during Tle3 secretion, we first validated the interactions of Tla3 with Tle3 and VgrG2b by a complementary approach of co-purification by affinity chromatography. Two different tagged versions of Tla3 were engineered by fusing a Strep-tag or a 10His-tag to the mature domain of Tla3 this leading to cytoplasmic tagged Tla3 proteins. The recombinant Tla3^C^_Strep_ was coproduced in *E. coli* BL21(DE3) pLysS with Tle3^C^_His_, and the Tla3^C^_His_ with 3 recombinant forms of Strep-tagged VgrG2b consisting in the full-length VgrG2b, or VgrG2b truncated for the extreme C-terminus (deletion of domain 3 in Fig. 3A) or VgrG2b truncated for the extreme C-terminus and the TTR domain (deletion of domains 2 and 3 in Fig. 3A). We initially tried with His-tagged VgrG2b but a problem of protein instability led us to shift for Strep-tagged VgrG2b. The bacterial lysates were loaded on a StrepTactin matrix, and Tla3^C^_Strep_ or the three recombinant VgrG2b_Strep_ were eluted with desthibiotin. The presence of Tle3^C^ _His_ and of Tla3^C^ _His_ was visualized in the elution fractions with anti-His antibodies. As shown in Fig. 4A, Tle3^C^ _His_ was found in the eluted fraction upon coproduction with Tla3^C^ _Strep_. We observed that Tla3^C^ _His_ is copurified only upon coproduction with the full length VgrG2b (Fig. 4B) or the VgrG2b truncated for the extreme C-terminus (Fig. 4C). Indeed when produced alone (Fig. 4B) or with VgrG2b truncated for the extreme C-terminus and the TTR domain (Fig. 4D), Tla3^C^ _His_ was not purified by affinity chromatography. Since VgrG2b truncated for the extreme C-terminus still copurified with Tla3 we can exclude that this domain is required for the interaction. This is in line with the BACTH assay that showed an interaction between Tla3 and a truncated VgrG2b consisting in only the DUF2345 and TTR domains (domains 1 and 2, Fig. 3A). Moreover, the deletion of the TTR domain affecting the copurification, one can conclude that this domain is key for the interaction. Taken together these data confirmed a direct interaction of Tla3 with Tle3 on one side and with VgrG2b on the other side. By taking into account the BACTH data and the copurification with two truncated forms of VgrG2b, the domain of interaction of VgrG2b with Tla3 can be delimitated to the TTR domain.

**Fig. 4:**
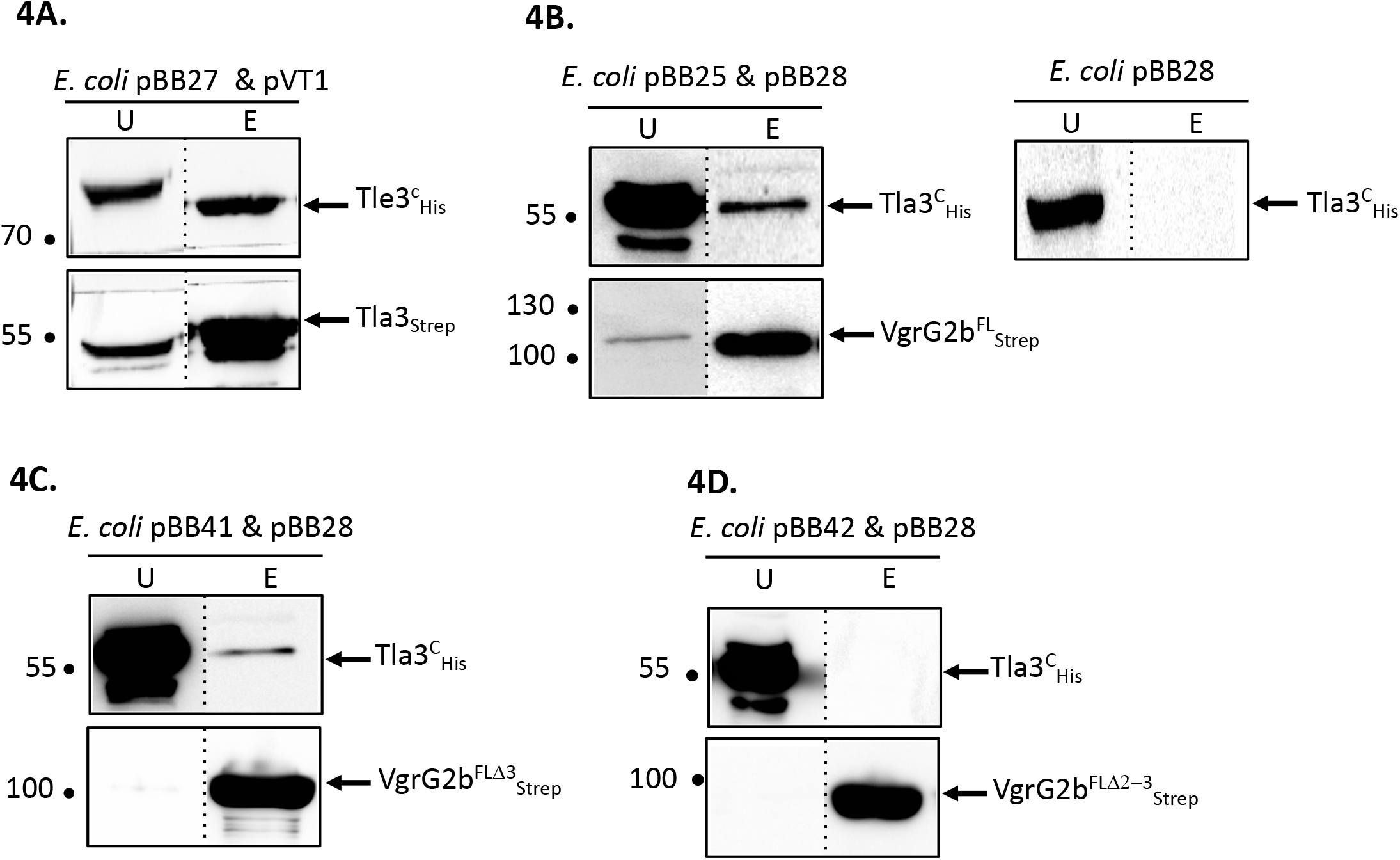
**Tla3 interaction with Tle3 (A) and VgrG2b (B-D)** Copurification assay on StrepTactin column of Tla3^c^_STREP_ with Tle3^c^_His_ produced *in E. coli* BL21(DE3)pLysS from pBB27 and pVT1 respectively (A) or of Tla3^c^_His_ with VgrG2b^FL^_STREP_ (B) or VgrG2b^FLD3^_STREP_ (C) or VgrG2b^FLD2-3^_STREP_ (D) produced from pBB28 and pBB25 (B) or pBB41(C) or pBB42 (D). The unbound (U) and eluted (E) fractions were collected and subjected to SDS-PAGE (10.5%) and Western blot analysis using anti-His antibody (Upper) and anti-streptavidin antibody (Lower). The dashed line separates lanes from non-adjacent part of the same gel. The position of the proteins and the molecular mass markers (in kDa) are indicated.

We then analyzed the cellular localization of Tla3 in *P. aeruginosa* that, according to its interactions with Tle3 and VgrG2b, should be cytoplasmic. As for Tli3, we engineered a chromosomally encoded Tla3_V5_ translational fusion (Supplementary Table 1). Tla3 was indeed immunodetected in the cytoplasmic fraction (Fig. 5A). One could note that in contrast with our first hypothesis suggesting an incorrect start codon for *tla3* and our observation of the recombinant protein in the periplasm of *E. coli* (Fig. 1 Sup), Tla3 was totally absent from the periplasmic fraction of *P. aeruginosa*. To strengthen this result, each putative ATG was individually mutated on the chromosome of the PAO1 strain encoding the Tla3_V5_ translational fusion (Fig. 5B). In agreement with its cytoplasmic localization in *P. aeruginosa*, Tla3 was only produced if the fourth ATG was intact. Accordingly the Tla3 protein synthesized from this ATG is not predicted to possess a N-terminal signal peptide (Fig. 1 Sup). In conclusion Tla3 is a cytoplasmic protein synthesized from the annotated translation start (pseudomonas.com) and this localization is in agreement with a role in the targeting of the toxin to the secretion machinery.

**Fig. 5:**
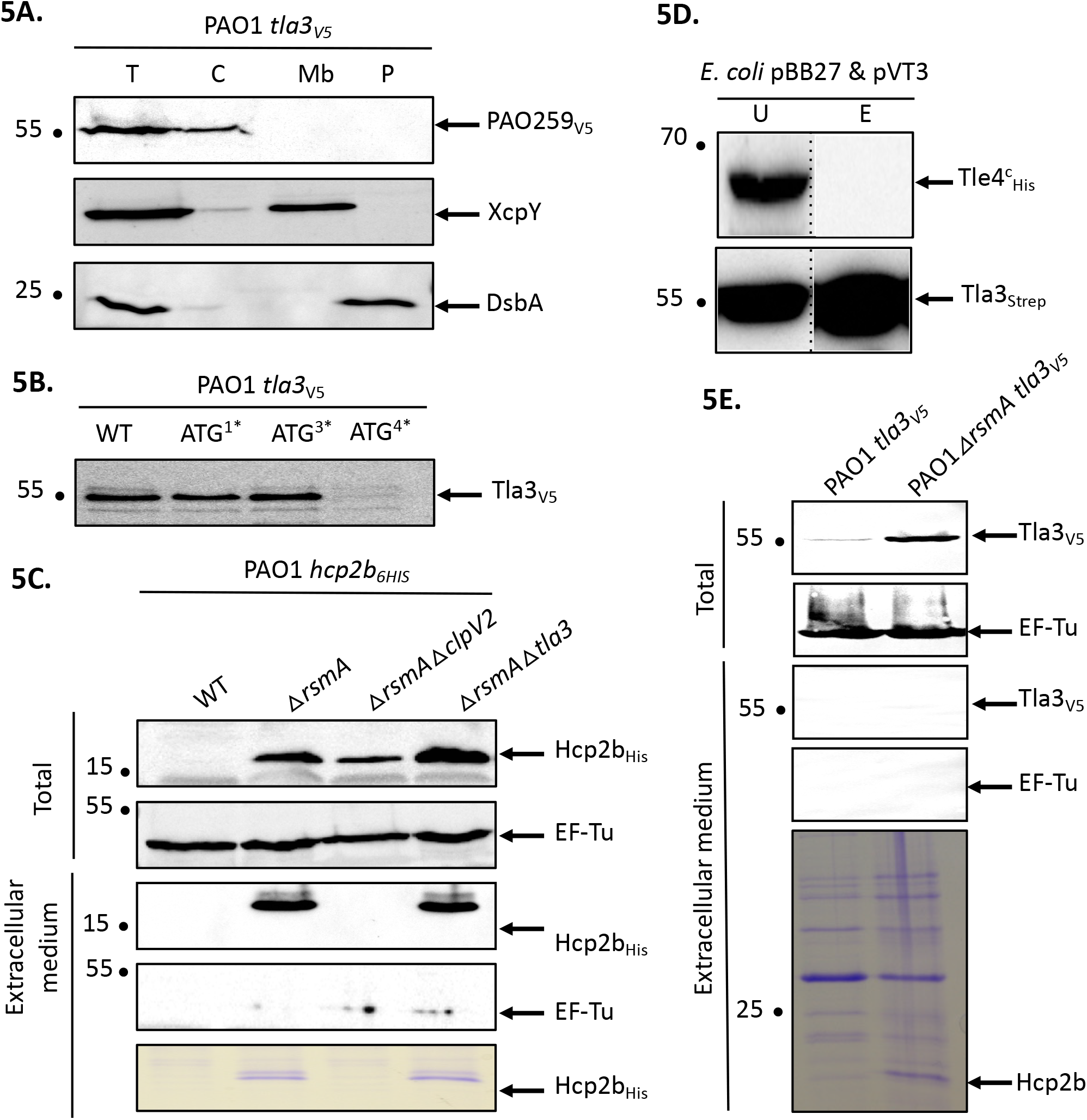
**Tla3(PA0259) is a cytoplasmic protein of *P. aeruginosa* (A)** Cells of *P. aeruginosa* PAO1 *tla3*_*V5*_ were subjected to fractionation and immunoblotting using antibodies directed against the V5 tag, XcpY and DsbA. XcpY and DsbA were used as membrane and periplasmic controls respectively. T: whole cell, C: cytoplasm, Mb: total membrane, P: periplasm. **The annotated ATG drives the initiation of translation of *tla3* in *P. aeruginosa* (B).** Immunodetection of Tla3_V5_ with anti-V5 antibodies produced in a WT background or in strains in which one of the four predicted ATG in *tla3* have been substituted in ATA. The number followed by a star indicates which ATG from Fig. 1Supp., ATG_4_ corresponding to the annotated ATG. **Tla3 is not required for Hcp2b secretion (C)**. Immunodetection of Hcp2b_6His_ with anti-His antibodies produced in a WT background (line 1) or in strains deleted for *rsmA* (lines 2 to 4) and *clpV2* (line 3) or *tla3* (line 4). The strains were grown at 25°C for 24 h and total bacteria were separated from extracellular medium. Anti-EF-Tu is used as a lysis control. The extracellular medium proteins were also stained with Coomassie-blue. **Tla3 doesn′t interact with Tle4 (D).** Copurification assay on StrepTactin column of Tla3^c^_STREP_ with Tle4_His_ produced *in E. coli* BL21(DE3)pLysS from pBB27 and pVT3 respectively. Legend as in Fig. 4. **Tla3 is not secreted (E**). Immunodetection of Tla3_V5_ with anti-HV5 antibodies produced in a WT background (line 1) or in strain deleted for *rsmA* (line 2). The strains were grown at 25°C for 24 h and total bacteria were separated from extracellular medium. Anti-EF-Tu is used as a lysis control. The extracellular medium proteins were also stained with Coomassie-blue. (A-E) The position of the proteins and the molecular mass markers (in kDa) are indicated.

Next, we asked whether Tla3 is specific for Tle3 or if it can be required for the secretion of other substrates of the H2-T6SS machinery. Since Hcp secretion is the hallmark of a functional secretion system, we studied the secretion of Hcp2b whose gene is upstream *vgrG2b* (Fig. 1A). Like Allsopp and colleagues (2017) we deleted the *rsmA* gene to enable Hcp2b production and thus secretion by a PAO1 strain encoding a Hcp2b_His_ translational fusion (Fig. 5C, compare line 1 and 2). RsmA is a posttranscriptional regulator known to repress all three T6SS clusters of *P. aeruginosa* [33]. This results in a massive secretion since Hcp2b_His_ can be observed in the extracellular protein samples by Coomassie-blue staining (Fig. 5C, lower panel). While Hcp2b_His_ secretion was abolished in a *rsmA clpV2* mutant, we observed that Hcp2b_His_ is still secreted in the absence of Tla3 (Fig. 5C, compare line 3 with line 4), suggesting that Tla3 is specific for the secretion of Tle3 but not for other H2-T6SS proteins. In line with this specific adaptor-toxin pair, Tla3^C^_Strep_ did not copurify TplE (Tle4), another antibacterial phospholipase delivered by the H2-T6SS machinery (Fig. 5D).

Finally as the interaction with a VgrG can suggest that Tla3 is itself a T6SS effector, we studied whether Tla3 is secreted by *P. aeruginosa*. To this end the *rsmA* gene was deleted from the PAO1 strain encoding a Tla3_V5_ translational fusion. Whereas Tla3_V5_ was better produced upon *rsmA* deletion (Fig. 5E), this did not lead to immunodetection of Tla3_V5_ in the extracellular medium although Hcp2b was readily observed by Coomassie-blue staining of the same samples. Tla3 is thus not an effector *per se*.

In conclusion the cytoplasmic localization of Tla3 in *P. aeruginosa* is appropriate with a recruiting role of Tle3 to the H2-T6SS machinery through an interaction with the TTR domain of VgrG2b. Tla3 is not co-secreted with Tle3. Tla3 role seems specific to Tle3 since it is not required for a functional H2-T6SS machinery and does not interact with TplE (Tle4), another H2-T6SS effector.

### Tle3 secretion mechanism

In order to study the antibacterial role and the secretion of Tle3 by *P. aeruginosa*, we performed intra-species bacterial competition assays between *P. aeruginosa* strains. This consists in studying the survival of prey bacteria lacking the *tli3* immunity gene (by CFU counting of antibiotic resistant bacteria) co-cultivated 24 hours at 37°C on plate with various attackers. As immunity genes are essential genes (protection from fratricide intoxication), both toxin and immunity genes have been deleted in order to construct a viable Δ*tli3*Δ*tle3* mutant strain (Supplementary Table 1). The figure 6 first confirms the antibacterial function of Tle3 and the immunity role of Tli3 since the growth of the immunity mutant was affected by the WT strain because it cannot resist Tle3 toxicity, while the Δ*tle3* mutant had no effect (Fig. 6 compare line 1 with line 2). The use of the Δ*clpV2* strain, a H2-T6SS mutant and of Δ*vgrG2b* and Δ*tla3* mutants allowed to demonstrate that Tle3 is delivered to prey bacteria through the H2-T6SS machinery and confirms that VgrG2b and Tla3 participate in Tle3 targeting to the H2-T6SS machinery (Fig. 6 compare line 1 with lines 4, 5 and 6). Indeed Δ*clpV2*, Δ*vgrG2b* and Δ*tla3* mutants had no effect on the immunity mutant growth since they cannot deliver Tle3 into prey bacteria. The complementation *in cis* of *tle3* and *tla3* deletions (the mutants have been constructed for this study) restored a WT phenotype (Fig. 6 compare line 2 with line 3, and line 6 with line 7) this demonstrating no polar effect on downstream genes. Furthermore the introduction of a wild-type copy of *tli3* at the *attB* site on *P. aeruginosa* chromosome of the Δ*tli3*Δ*tle3* mutant restores wilt-type competition capacity to this strain (Fig. 4A Sup). This confirms that the absence of the immunity was responsible of the phenotypes observed for the Δ*tli3*Δ*tle3* strain (Fig. 6). Finally, the Δ*tla3* mutant has been used as a prey and was not affected by the WT strain or any of the mutants excluding definitively a role of immunity as proposed by its annotation (Pseudomonas.com) and confirming its adaptor function (Fig. 4B Sup).

**Fig. 6:**
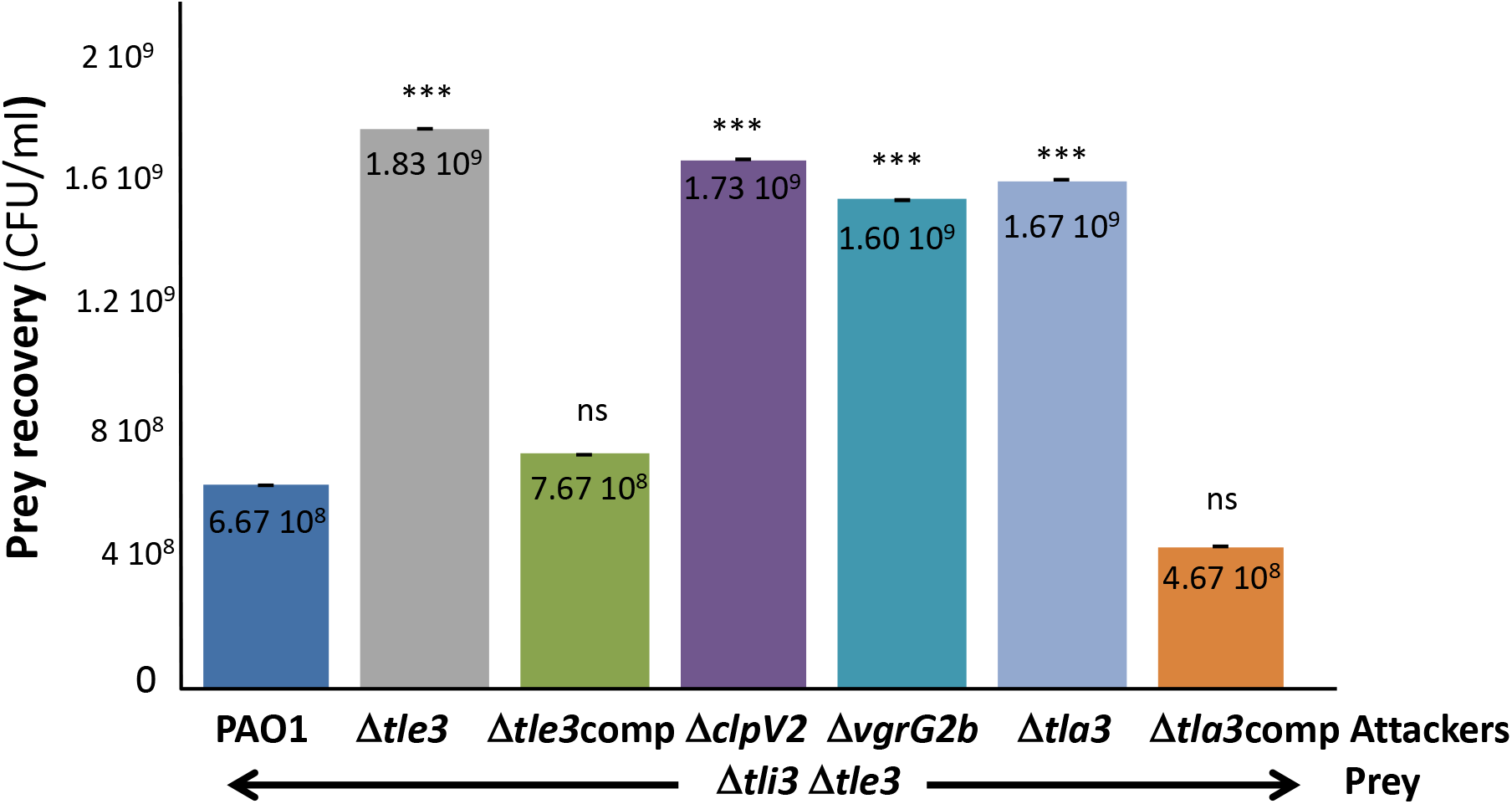
***P. aeruginosa* growth competition.** The *P. aeruginosa* prey strain (Δ*tli3*Δ*tle3*) was incubated with various *P. aeruginosa* attacker strains as indicated in the figure for 24 h at 37°C. The number of recovered prey bacteria is indicated in CFU/ml. “comp” stand for *cis* complementation of the corresponding mutation with a wild-type copy inserted at the *attB* site on *P. aeruginosa* chromosome. Error bars represent ± SEM (n = 3); ***p < 0.001, ns not significant.

Taken together these results demonstrated that Tle3 is an effective H2-T6SS-dependent antibacterial toxin loaded onto the VgrG2b puncturing device via Tla3 and neutralized by Tli3 in resistant prey bacteria.

### VgrG2b is a trans-kingdom toxin

A putative neutral zinc metallopeptidase domain has been predicted at the extreme C-terminus of VgrG2b by Pukatzki and colleagues [34] (Fig. 3A). This motif (Prosite PS00142, PFAM04298) consists in a metal-binding consensus motif HExxH, the two histidine residues being ligands of the catalytic Zn^2+^ and the glutamic acid residue involved in nucleophilic attack. As an effector with a protease activity can target both eukaryotic and bacterial proteins, we searched for an antibacterial activity of the VgrG2b C-terminal extension. To do this we performed the same heterologous toxicity assay in *E. coli* as with Tle3. As shown in figure 7, whereas the production of VgrG2b_Cter_ in the cytoplasm did not impact *E. coli* growth, its periplasmic production killed *E. coli* (Fig. 7, compare line 2 and 5). Moreover substitution of the histidine in position 935 and of the glutamic acid 936 for an alanine relieves VgrG2b_Cter_ toxicity this showing that VgrG2b is a novel antibacterial protease active in the periplasm.

**Fig. 7:**
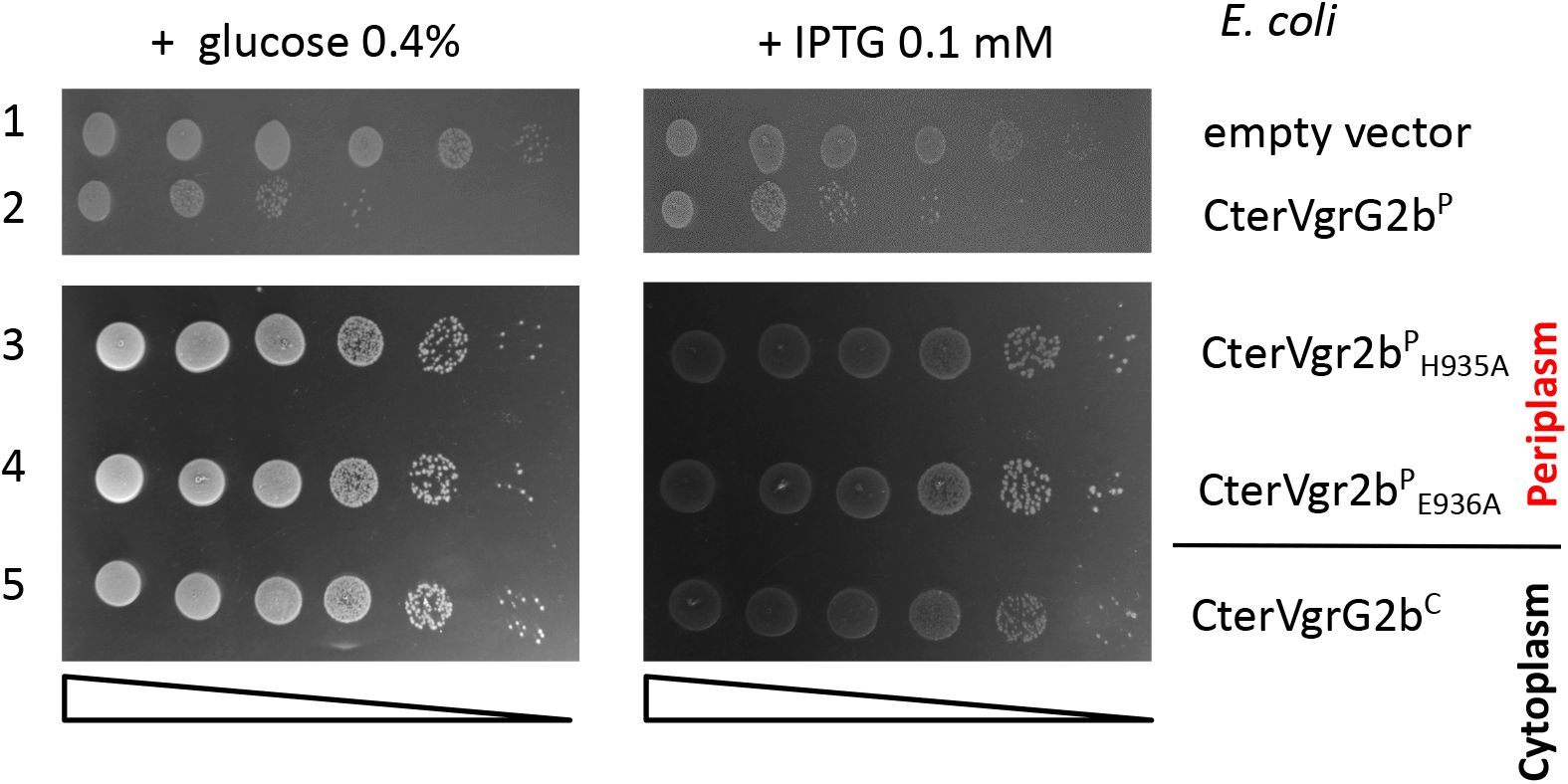
**The protease domain of VgrG2b is required for its antibacterial activity.** Serial dilutions (from non-diluted to 10^−5^) of normalized cultures of *E. coli* BL21(DE3)pLys producing the wild-type C-terminal domain of VgrG2b in the cytoplasm, called CterVgrG2b^C^(from pBB43, a pETDuet-1 derivate, line 5) or in the periplasm, called CterVgrG2b^P^ (from pBB44, a pET22b(+) derivate yielding a fusion of the C-terminal domain with a Sec signal peptide, line 2) or two variants CterVgr2b^P^ _H935A_(from pBB45, line 3) and CterVgr2b^P^ _E936A_ (from pBB46, line 4) were spotted on LB agar plates supplemented (left panel) with 0.4% glucose or (right panel) with 0.1mM IPTG. Glucose and IPTG allow respectively repression and induction of the gene encoding the T7 RNA polymerase.

## Discussion

Here we report the existence of a novel pair of antibacterial effector and immunity of the H2-T6SS of *P. aeruginosa*, Tle3 (PA0260) and Tli3 (PA0261), and we propose a chronology of Tle3 secretion process that includes a cytoplasmic adaptor protein, Tla3 (PA0259) to load the toxin onto the VgrG2b spike (a model is proposed in Fig. 8). Through heterologous toxicity assay and bacterial competition, we show that Tle3 was toxic once delivered in the periplasm of prey bacteria and that Tli3 can neutralize the toxin in this compartment. Interestingly this led us to discover that VgrG2b that we previously recognized as an anti-eukaryotic effector possesses an antibacterial activity as well.

**Fig. 8:**
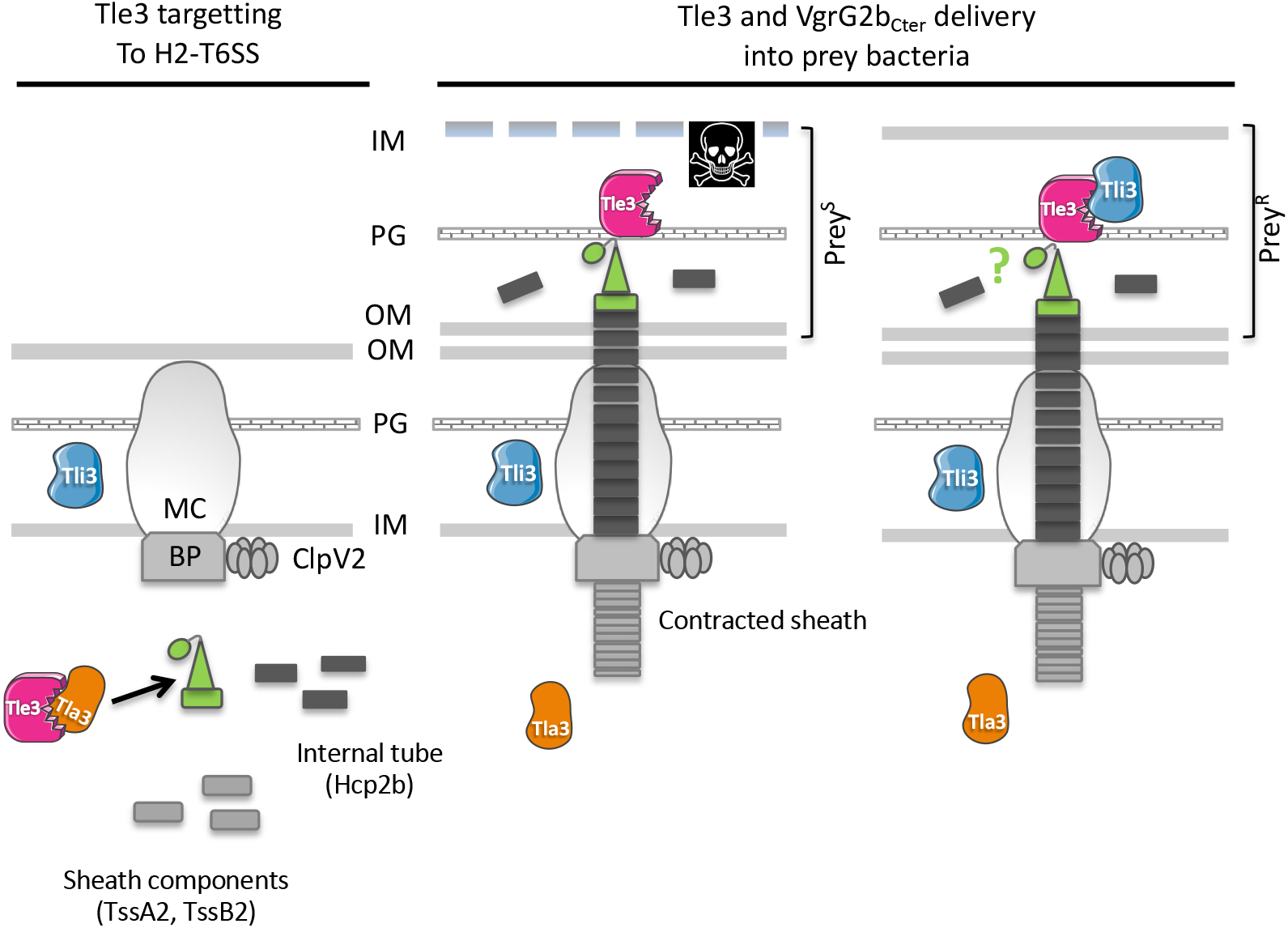
**Working model: Tle3 targeting to the H2-T6SS machinery (left panel)** Tle3 (in pink) is taken in charge in the cytoplasm by Tla3 (in orange) that binds VgrG2b (green). Upon Hcp2b (in dark grey) assembly into the growing sheath (light grey), the VgrG2b loaded with the Tle3 is placed at the tip of the Hcp arrow. **Tle3 and VgrG2b_Cter_ delivery into bacteria (right panel).** The sheath contraction in the cytoplasm propels the Hcp arrow towards the target bacterium. Tle3 associated with this expelled structure is thus translocated into target cells, as well as VgrG2b_Cter_ (green circle). The prey is killed (Prey^S^) except if it has the Tli3 immunity (in blue) in the periplasm (Prey^R^). The mechanism of resistance to VgrG2b_Cter_ is still unknown. OM: outer membrane, PG: peptidoglycan, IM: inner membrane, BP: base plate, MC: membrane complex. Prey^S^: sensitive prey, Prey^R^: resistant prey.

The VgrG-recruitment of cargo effectors has been previously evidenced for several antibacterial effectors among them two toxins of the Tle family, TseL (Tle2) of *V. cholerae* [16,17,35] and Tle1 of EAEC [29]. In both cases the *tle* genes were just downstream the *vgrG* genes like the organization of *vgrG2b* and *tle3* of *P. aeruginosa*. TseL and Tle1 have been shown to directly bind a dedicated VgrG, VgrG3 and VgrG1 respectively [29,35]. The domain of interaction within EAEC VgrG1 has been delimitated to the TTR domain and may also include the DUF2345, both of which are present in the *P. aeruginosa* VgrG2b. In line with this, and despite Tle3 requiring an adaptor to be targeted to VgrG2b, we have shown that Tla3 interacts with the TTR domain of VgrG2b. Taken together these data demonstrate that TTR domains of VgrGs are involved in recruitment and transport of Tle effectors, directly or through adaptor. Likewise C-terminal extensions of VgrG1 and VgrG2 of *Agrobacterium tumefaciens* were identified as specifically required for the delivery of each cognate DNAse toxins, named Tde1 and Tde2 respectively [22]. C-terminal domains of VgrGs can thus be considered more generally as specificity determinants for T6SS effector loading and transport.

Interestingly TseL of *V. cholerae* requires also Tap-1 (Tec) as an adaptor protein to be delivered to another VgrG, called VgrG1 [16,17], showing that a sole toxin can be targeted directly and indirectly to two different VgrG proteins. Tap-1 (Tec) belongs to the DUF4123 family of adaptor proteins that contains also VasW of *V. cholerae* [18] and several uncharacterized gene products linked to effector genes with a MIX (marker for type VI effectors) motif in *Proteus mirabilis* or *B. thailandensis* for instance [19]. Interestingly TecT, a DUF4123 adaptor of *P. aeruginosa*, has been shown to require a co-adaptor, called co-TecT, to deliver the TseT effector to the PAAR4 protein [27]. This is the first example of an adaptor-co-adaptor module. Taken together these data suggest a conserved role for DUF4123 adaptors in the recruitment of a number of T6SS effectors. Remarkably Tla3 of *P. aeruginosa* does not belong to the DUF4123 adaptor family, or to that of the two other unrelated families, the DUF1795 adaptor family, reported with EagR (effector-associated gene with Rhs) in *Serratia marcescens* [21] or EagT6 in *P. aeruginosa* [20], and the DUF2169 adaptor family reported with Atu3641 in *A. tumefaciens* [22]. Nor Tla3 is a PAAR protein, the last class of effector targeting mode to a VgrG [22,27,36–38]. Instead we find that Tla3 harbors two DUF2875 domains (Fig. 3A) that are both required for the interaction with the toxin. Moreover genes coding DUF2875 containing proteins can be find at the vicinity of *tle, tli, vgrG* or PAAR genes, but they are restricted to α and β proteobacteria (Fig. Sup 5). We thus hypothesize that DUF2875 might assist in T6SS-mediated effector delivery. Like the three other adaptor families (DUF4123, DUF1795, DUF2169), we have observed that Tla3 is not required for the H2-T6SS functionality since the Δ*tla3* mutant still secrete Hcp2b and can compete with a WT strain, and that Tla3 is specific for the Tle3 toxin since it did not interact with another H2-T6SS effector, TplE (Tle4). Finally as we did not detect Tla3 secretion under constitutive H2-T6SS condition, we propose that Tla3 hands over Tle3 to VgrG2b in the cytoplasm prior to its loading to the baseplate and further recruitment to the central Hcp tube in preparation (Fig. 8). Overall, the existence of various modes of effector recruitment, further refined with adaptors, likely explains how the T6SS is able to deliver numerous and structurally diverse proteins.

Five families of Tle, Tle1-5, have been identified among Gram-negative bacteria [11] and four Tle have been studied in *P. aeruginosa* so far. Our demonstration of the activity of Tle3 in the periplasm is consistent with the observations that the heterologous periplasmic production of PldA (Tle5a) [11,12], PldB (Tle5b) [12], Tle1 [39] and TplE (Tle4) [13] is toxic. The reason of the periplasmic activity of Tle proteins is still unclear although several hypothesis have been proposed [29], the most likely being an activation of the toxin in this compartment. Very recently this has nicely been exemplified with the hijacking of DsbA in the target cells of *Serratia macescens* for the activation of incoming effectors [40]. No member of the Tle3 family has yet been enzymatically characterized. Our attempts to efficiently purify Tle3 from *E. coli* or from *P. aeruginosa* have been unsuccessful, even if we have noticed that the presence of the Tla3 adaptor stabilized Tle3, it still formed inclusion bodies. In the future we will decipher the enzymatic activity of Tle3, which is presumably active on membrane phospholipids as our preliminary data of thin-layer chromotography tend to show.

The periplasmic activity of Tle toxins is counteracted by the synthesis of a cognate immunity protein that is usually a periplasmic soluble protein, as we showed for Tli3 in *P. aeruginosa*, or a membrane-anchored lipoprotein [11]. Interestingly the genetic organization of the *tli3* gene upstream of the *tle3* gene observed in *E. coli, K. pneumoniae, B. cenocepacia* or *R. solanacearum* [11] is conserved in *P. aeruginosa*. The fact that the two genes are co-transcribed (the immunity being the first) is key for the protection against toxicity. Indeed we have observed systematic protection against the periplasmic toxicity of Tle3 in *E. coli* when the two genes were expressed from the same promoter under the same plasmid whereas it was not as efficient when the genes were on two plasmids. This genetic link reinforces the connection within the toxin/immunity pair. This has previously been noticed with a T7SS antibacterial toxin and its immunity in *Staphylococcus aureus* [41]. Other immunities of Tle characterized so far have been shown to inhibit the action of the effector by direct protein-protein contacts [11,12,29]. Our copurification assay in *E. coli* demonstrates a direct interaction between Tle3 and Tli3 that was already suggested with the release of Tle3 toxicity upon coproduction of Tli3 in the periplasm. A crystal structure of the *P. aeruginosa* TplE (Tle4) effector in complex with its immunity protein TplEI (Tli4) revealed that the immunity uses a grasp mechanism to prevent the interfacial activation of the toxin [30].

Like two other H2-T6SS related orphan *vgrG* loci, the *vgrG4b* cluster encoding PldA (Tle5a) and the *vgrG2a* encoding TplE (Tle4), we show here that the *vgrG2b* cluster has both antibacterial activities (through Tle3 and VgrG2b) and anti-eukaryotic (through VgrG2b; [26]). Interestingly VgrG2b is thus (i) a structural component of the H2-T6SS puncturing device since our bacterial competition showed its requirement for Tle3 delivery, (ii) an anti-eukaryotic effector through an interaction with the microtubule nucleating complex [26] and (iii) an antibacterial effector as suggested by our toxicity assay in *E.coli*. We have shown that two conserved residues of the putative metallopeptidase motif (an histidine and a glutamic acid) are essential for the VgrG2b antibacterial activity. This is consistent with an antibacterial protease activity for VgrG2b that will be, to our knowledge, the first case in the T6SS effector literature. The discovery of the VgrG2b immunity is even more exciting. Could it share Tli3 with Tle3? We also hypothesize that an auto-immunity mechanism could exist in which an immunity domain within VgrG2b could be released upon auto-processing liberating the active protease domain like the Serine protease autotransporter proteins.

## Materials and Methods

### Bacterial strains, growth conditions and plasmid construction

All *P. aeruginosa* and *E. coli* strains used in this study are described in Supplementary Table 1. Briefly, the *E. coli* K-12 DH5α and CC118λPir were used for cloning procedures. The BL21(DE3)pLysS and BTH101 were used for protein production and BACTH analyses respectively. Strains were grown in LB or in TSB medium (for *P. aeruginosa*) at 37°C or 30°C. Specific growth conditions are specified in the text when necessary. Recombinant plasmids were introduced into *P. aeruginosa* by triparental mating using the conjugative properties of the helper plasmid pRK2013 (Supplementary Table 1). Plasmids were maintained by the addition of ampicillin (50 μg/mL), kanamycin (50 μg/mL), chloramphenicol (30 μg/mL), streptomycin (30 μg/mL for *E. coli*, 2000 μg/mL for *P. aeruginosa*) or gentamicin (30 μg/mL for *E. coli*, 115 μg/mL for *P. aeruginosa*). Expression of genes from pT7 in BL21(DE3)pLysS was blocked with 0.4% of glucose and induced in exponential phase (OD_600_=0.4–0.6) for 3 hours with 1mM of IPTG. Cloning procedures were described in [26]. The plasmids used and constructed are described in Supplementary Table 1, the list of oligonucleotides (synthesized by Eurogentec or IDT) is given in Supplementary Table 2.

### Cloning procedures for ***P. aeruginosa* mutants**

To generate *P. aeruginosa* mutants, 500 bp upstream and 500 bp downstream of the gene to be deleted were amplified by overlapping PCR with Q5 high fidelity DNA polymerase (NEB) using primers listed in Supplementary Table 1. The PCR product was cloned in pKNG101 suicide vector by one-step sequence and ligation-independent cloning (SLIC) [42], which was then sequenced. pKNG101 derivatives, maintained in the *E. coli* CC118λpir strain, were mobilized in *P. aeruginosa* strains. The mutants, in which the double recombination events occurred, were confirmed by PCR analysis.

### Heterologous toxicity assays

*E. coli* BL21(DE3)pLysS containing plasmids producing cytoplasmic or periplasmic targeted proteins were grown overnight at 37°C in LB with 0.4% of glucose. 10 μL drops of bacterial suspensions serially diluted were spotted onto LB agar plates containing 0.1 mM IPTG or 0.4 % glucose and cells were grown for 16 h at 37°C.

### Bacterial two-Hybrid Assay

Protein-protein interactions were assessed with the adenylate cyclase-based two-hybrid technique using protocols published previously [43,44]. Briefly, the proteins to be tested were fused to the isolated T18 and T25 catalytic domains of the *Bordetella* adenylate cyclase. After introduction of the two plasmids producing the fusion proteins into the reporter BTH101 strain, plates were incubated at 30°C for 24 h. Three independent colonies for each transformation were inoculated into 600 μl of LB medium supplemented with ampicillin (50 μg/mL), kanamycin (50 μg/mL), and IPTG (0.5 mM). After overnight growth at 30 °C, 5 μl of each culture was spotted onto LB agar plates supplemented with ampicillin, kanamycin, IPTG, and 5-bromo-4-chloro-3-indonyl-D-galactopyrannoside (X-gal, 40μg/mL) and incubated for 16 h at 30 °C.

### Protein purification by affinity chromatography

*Escherichia coli* BL21(DE3)plysS cells carrying the pRSFDUET-1 and pETDUET-1 derivates were grown at 37°C in LB to an OD_600_ *∼* 0.5 and the expression of the PA0262, PA0261, PA0260 or PA0259 genes was induced with IPTG (1 mM) for 3 h at 37°C. Cells were harvested by centrifugation at 1914 × g for 30 min at 4°C. The cell pellet was resuspended in Tris-HCl 50 mM pH 8.0, NaCl 150 mM, Triton X-100 0.1%, lysozyme 0.5 mg/mL and EDTA 1 mM and stored at −80°C. Cells were supplemented with DNase (20 μg/mL), MgCl_2_ and phenylmethylsulfonyl fluoride 1mM and cells were lysed by three passages at the Emulsiflex-C5 (Avestin), and lysates were clarified by centrifugation at 16000 × g for 30 min. The supernatant was loaded onto a 5-mL StrepTrap HP (GE Healthcare) column and then washed with 50 mM Tris-HCl pH 8.0, 150 mM NaCl at 4°C. The fusion protein was eluted in the affinity buffer supplemented with 2.5 mM desthibiotin. Peak fractions were pooled and loaded onto a Superose 200 10/300 column (GE Healthcare) equilibrated in 50 mM Tris-HCl pH 8.0, 50 mM NaCl.

### Fractionation of ***P. aeruginosa***

Fractionation of cells into spheroplasts (cytoplasm and membranes) and periplasmic fractions were done as previously described [45]. Proteins corresponding to the cytoplasm and periplasm fractions or to insoluble material were resuspended in loading buffer.

### Protein secretion

*P. aeruginosa* strains were grown at 25°C in TBS for 24 hours. Cells corresponding to 10 units DO_600_ and extracellular medium were separated by centrifugation at 2000 × g for 10 min at room temperature. 2/3 of the supernatants were collected and centrifuged at 13 000 × g for 5 min at room temperature. Proteins contained in the supernatant were precipitated with tricholoro-acetic acid (TCA, 15%) for 3 h at 4°C. Samples were centrifuged at 13 000 × g for 30 min at 4°C, pellets washed with 90% acetone and resuspended in loading buffer.

### SDS-PAGE and western-blot

Protein samples derived from equivalent amounts of culture (i.e. optical density equivalents) resuspended in loading buffer were boiled and separated by SDS-PAGE. Proteins were then stained by Coomassie-blue or immunodetected as described before [26] using primary polyclonal antibodies directed against His6 epitope-tag (Penta His, Qiagen, dilution 1:1000), V5 epitope-tag (Bethyl Laboratories, dilution 1:1000), Strep epitope-tag (IBA StrepMAB Classic, dilution 1:000), DsbA (kindly gifted by K.E. Jaeger – university of Heinrich-Heine, dilution 1:25000), or monoclonal antibodies directed against EF-Tu (Hycult-biotech, dilution 1:20000), XcpY (laboratory collection, dilution 1:5000), TolB (laboratory collection, dilution 1/500). Peroxidase-conjugated anti-Mouse or anti-Rabbit IgGs (Sigma, dilution 1:5000) were used as secondary antibodies. Nitrocellulose membranes were revealed with homemade enhanced chemiluminescence and were scanned using ImageQuant LAS4000 analysis software (GE Healthcare Life sciences).

Protein samples equivalent to 0.1 OD_600_ units were loaded for whole cell and spheroplasts analysis while protein samples equivalent to 0.2 OD_600_ units were used for cytoplasm or periplasm analysis and protein samples equivalent to 1 OD_600_ units were used for extracellular medium analysis.

### Bacterial Competition assays

Intraspecific competition assays between *P. aeruginosa* strains were performed as previously described [12] with modifications. The prey cells carry pJN105 vector (Gm^R^) to allow counterselection. Overnight cultures of *P. aeruginosa* attacker and prey cells were mixed in a 5 : 1 (attacker : prey) ratio and harvested by centrifugation at 3724 × g for 5 min. The pellet was resuspended in 200 μL of PBS 1X and spotted onto 0.45-μm nitrocellulose membranes overlaid on a 1 % bactoagar plate. After 24 hours of incubation at 37°C, cells were resuspended in 2 mL of PBS 1X, normalized to an OD_600nm_ of 0.5 and 10 μL of bacterial serially diluted (10^−1^ to 10^−6^) were spotted onto selective LB agar plates containing gentamicin (125 μg/mL). Significant growth difference of the prey bacteria for each competition assay was computed by one-way ANOVA (Stat Plus) and unpaired Student’s Test (Excel).

## Acknowledgments

We thank V. Tutagata for pVT1, pVT8 and pVT9 constructs, B. Douzy for all the advices during protein purification and members of B.B. PhD committee for helpful discussion and support. We are grateful to M. Ba, I. Bringer, A. Brun and O. Uderso for technical assistance. B.B. was financed with a PhD fellowship from the French Research Ministry. This work was supported by recurrent funding from the CNRS and Aix-Marseille University. The project leading to this publication has received funding from the Excellence Initiative of Aix-Marseille University-A*Midex, a French “Investissements d′Avenir” program (“Emergence & Innovation” A-M-AAP-EI-17-139-170301-10.31-BLEVES-HLS).

## Author contributions

B.B. and S.B. designed and conceived the experiments. B.B., C.S., and S.D. performed the experiments. S.B. supervised the execution of the experiments. B.B., C.S., B.I. and S.B. analyzed and discussed the data. S.B. wrote the paper with contribution from B.B. and reading from B.I.

## Abbreviations

T6SS: Type VI secretion system
Tle: Type VI lipase effector
Tli: Type VI lipase immunity
1914Tla: Type VI lipase adaptor

