## Supplementary data for "AT6SS trans-kingdom effector is required for the delivery of a novel antibacterial toxin in *Pseudomonas aeruginosa*"

**Supplementary figure 1:** *In silico* analysis of the nucleic and protein sequence of Tla3 (PA0259).

**Supplementary figure 2:** Production of Tle3$^{p}$ (S2A) and localization of Tle3$^{c}$, Til3 and Tla3$_{ATG1}$ in *E. coli* BL21(DE3)pLysS (S2B-D).

**Supplementary figure 3:** Bacterial two-hybrid assay.

**Supplementary figure 4:** *P. aeruginosa* growth competition.

**Supplementary figure 5:** Adaptors of Tla3 family DUF2875 are restricted to beta and gamma proteobacteria.

**Table 1:** *E. coli* and *P. aeruginosa* strains and plasmids used in this study.

| | Genotype or description | Origin |
|---|---|---|
| **Strains** | | |
| ***E. coli*** | | |
| DH5α | F-, Δ(*argF -lac* )U169, *phoA* , *supE44* , Δ(lacZ)M15, *relA* , *endA* , *thi* , *hsdR* | Invitrogen |
| CC118λpir | Δ(*ara-leu* ) *ara* D Δ*lac* X74 *gal* E *gal* K *pho* A20 *thi* -1 *rps* E *rpo* B *arg* E (Am) *rec* A1 RfR (λpir) | Laboratory collection |
| BL21(DE3)PlysS | F– *ompT gal dcm lon hsdSB(rB–mB–)* λ(DE3 [*lacI lacUV5-T7p07 ind1 sam7 nin5* ]) [*malB+]K-1* 2(λS) pLysS[T7p20 orip15A, CmR | Laboratory collection |
| BTH101 | F-, *cya-99, araD139, galE15, galK16, rpsL1 (Str r), hsdR2, mcrA1, mcrB1.* | Laboratory collection |
| ***P. aeruginosa*** | | |
| PAO1 | Wild-type prototroph, cytotoxic and invasive | Laboratory collection |
| PAO1 *tla3* V5 | chromosomally encoded Tla3V5 translational fusion in PAO1 | This work |
| PAO1 △*rsmA* | *rsmA* deletion mutant | This work |
| PAO1 *hcp2b 6His* | chromosomally encoded H*cp2b 6His* translational fusion in PAO1 | This work |
| PAO1 △*rsmA hcp2b His* | chromosomally encoded *hcpC6His* translational fusion in PAO1 △*rsmA* | This work |
| PAO1 △*rsmA* △*clpV2 hcp2b His* | chromosomally encoded *hcpC6His* translational fusion in PAO1 △*rsmA* △*clpV2* | This work |
| PAO1 △*rsmA* △*tle3 hcp2b His* | chromosomally encoded *hcpC6His* translational fusion in PAO1 △*rsmA* △*tle3* | This work |
| PAO1 △*rsmA* △*tla3 hcp2b His* | chromosomally encoded *hcpC6His* translational fusion in PAO1 △*rsmA* △*tla3 (PA0259)* | This work |
| PAO1 tli3V5 | chromosomally encoded Tli3V5(PA0261) translational fusion in PAO1 | This work |
| PAO1 *tla3V5* | chromosomally encoded Tla3V5(PA0259) translational fusion in PAO1 | This work |
| PAO1△*clpV2* | *clpV2* deletion mutant | [1] |
| PAO1△*vgrG2b* | *vgrG2b* deletion mutant | [2] |
| PAO1△*tli3* △*tle3* | *tli3 (PA0261)* and *tle3* deletion mutant | This work |
| PAO1△*tle3* | *tle3* deletion mutant | This work |
| PAO1△*tla3* | *tla3 (PA0259)* deletion mutant | This work |
| PAO1△*tli3* △*tle3 attB :: tli3+* | *cis* complementation of △*tli3* △tle3 mutation by a *tli3* copy inserted at the *attB* site on *P. aeruginos* a chromosome | This work |
| PAO1△*tle3* attB :: *tle3* + | *cis* complementation of △*tle3* mutation by a *tle3* copy inserted at the *attB* site on *P. aeruginosa* chromosome | This work |
| PAO1△*tla3* attB :: *PA0259* + | *cis* complementation of △*tla3* mutation by a *tla3* copy inserted at the *attB* site on *P. aeruginosa* chromosome | This work |
| **Plasmids** | | |
| pRK2013 | ColE1 origin, Tra+, mob+, KmR | Laboratory collection |
| pKNG101 | R6K origin, TRK2 origin, mobRK2, sacBR+, SmR | [3] |
| pKNG101△*rsmA* | suicide vector for *rsmA* deletion, ,SmR | [4] |
| pBB29 | suicide vector for *tle3* deletion, SmR | This work |
| pBB30 | suicide vector for *tla3* (PA0259) deletion, SmR | This work |
| pBB31 | suicide vector for *tli3 (PA0261)* and *tle3* deletion, SmR | This work |
| pBB32 | suicide vector for *tli3* V5 insertion, SmR | This work |
| pBB33 | suicide vector for *tla3* V5 insertion, SmR | This work |
| pBB34 | suicide vector for *hcp2b* 6His insertion, SmR | This work |
| pBB35 | suicide vector for ATG1 substitution in ATA upstream the annotated ATG of *tla3* *(PA0259)* , SmR | This work |
| pBB36 | suicide vector for ATG3 substitution in ATA upstream the annotated ATG of *tla3 (PA0259)* , SmR | This work |
| pBB37 | suicide vector for annotated ATG substitution in ATA of *tla3 (PA0259)* , SmR | This work |
| Mini-CTX1 | Contains *attP* site for integration at the *attB* site of *P. aerugionosa* chromosome, TcR | [5] |
| pBB38 | *tle3* under control of *hcp2b* promoter cloned in EcoRI of Mini-CTX1, TcR | This work |
| pBB39 | *tli3 (PA0261)* under control of *hcp2b* promoter cloned in EcoRI of Mini-CTX1, TcR | This work |
| pBB40 | *tla3 (PA0259)* under control of *hcp2b* promoter cloned in EcoRI of Mini-CTX1, TcR | This work |
| pJN105 | *P. aeruginosa* expression vector with an arabinose inducible pBAD promoter, GmR | [6] |
| pUT18 | Bacterial Two Hybrid vector, ColE1 origin, P*lac* , T18 fragment of *Bordetella pertussis* CyaA, AmpR | Euromedex |
| pBB1 | *tle3* cloned downstream the T18 coding sequence in pUT18 | This study |
| pBB2 | *tle3(4-380)* cloned dowstream the T18 coding sequence in pUT18 | This study |
| pBB3 | *tle3(384-717)* cloned dowstream the T18 coding sequence in pUT18 | This study |
| pBB4 | *PA0261* cloned downstream the T18 coding sequence in pUT18 | This study |
| pBB5 | *PA0259* cloned downstream the T18 coding sequence in pUT18 | This study |
| pBB6 | *PA0259(21bp upstream ATG3-244)* cloned downstream the T18 coding sequence in pUT18 | This study |
| pBB7 | *PA0259(247-381)* cloned downstream the T18 coding sequence in pUT18 | This study |
| pBB8 | *PA0262* (1-2) cloned downstream the T18 coding sequence in pUT18 | This study |
| pBB9 | *PA0262* (1-2-3) cloned downstream the T18 coding sequence in pUT18 | This study |
| T18-Pal | *pal* cloned downstream the T18 coding sequence in pUT18 | [7] |
| pKNT25 | Bacterial Two Hybrid vector, p15A origin, P*lac* , T25 fragment of *Bordetella pertussis* CyaA, KanR | Euromedex |
| pBB10 | *tle3* cloned downstream the T25 coding sequence in pkNT25 | This study |
| pBB11 | *PA0261* cloned downstream the T25 coding sequence in pkNT25 | This study |
| pBB12 | *PA0259* cloned downstream the T25 coding sequence in pkNT25 | This study |
| pBB13 | *PA0262* (1-2) cloned downstream the T25 coding sequence in pkNT25 | This study |
| pBB14 | *PA0262* (1-2-3) cloned downstream the T25 coding sequence in pkNT25 | This study |
| pUT18C | Bacterial Two Hybrid vector, ColE1 origin, P*lac* , T18 fragment of *Bordetella pertussis* CyaA, AmpR | Euromedex |
| pBB15 | *tle3* cloned upstream the T18 coding sequence in pUT18C | This study |
| pBB16 | *PA0261* cloned upstream the T18 coding sequence in pUT18C | This study |
| pBB17 | *PA0259* cloned upstream the T18 coding sequence in pUT18C | This study |
| pBB18 | *PA0262* (1-2) cloned usptream the T18 coding sequence in pUT18C | This study |
| pBB19 | *PA0262(1-2-3)* cloned upsteam the T18 coding sequence in pUT18C | This study |
| pKT25 | Bacterial Two Hybrid vector, p15A origin, P*lac* , T25 fragment of *Bordetella pertussis* CyaA, KanR | Euromedex |
| pBB20 | *tle3* cloned upstream the T25 coding sequence in pKT25 | This study |
| pBB21 | *PA0261* cloned upstream the T25 coding sequence in pKT25 | This study |
| pBB22 | *PA0259* cloned upstream the T25 coding sequence in pKT25 | This study |
| pBB23 | *PA0262* (1-2) cloned usptream the T25 coding sequence in pKT25 | This study |
| pBB24 | *PA0262(1-2-3)* cloned upsteam the T25 coding sequence in pKT25 | This study |
| ToLB-T25 | *tolB* cloned upstream the T25 coding sequence in pkT25 | [7] |
| pRSFDUET-1 | Expression vector, RSF origin, *lacI* , PT7, KanR | Novagen |
| pBB25 | *vgrG2b* cloned into pRSFDUET-1, C-terminal Strep epitope, KanR | This study |
| pBB26 | *PA0261* downstream the signal sequence cloned into pRSFDUET-1, C-terminal Strep epitope, KanR | This study |
| pVT8 | *PA0261* cloned into pRSFDUET-1, C-terminal V5 epitope, KanR | This study |
| pBB27 | *PA0259* from ATG3 cloned into pRSFDUET-1, C-terminal Strep epitope, KanR | This study |
| pVT9 | *PA0259* from ATG1 cloned into pRSFDUET-1, C-terminal V5 epitope, KanR | This study |
| pSBC107 | insert from pSBC81 cloned into pVT8 leading to *tli3* and the sequence coding Tle3P on the same transcript, KanR | This study |
| pETDuet-1 | Expression vector, ColE1 origin, *lacI* , *PT7* , ApR | Novagen |
| pVT1 | *tle3* cloned into pETDUET-1, C-terminal 10His epitope, ApR | This study |
| pBB28 | *tla3 (PA0259)* from *ATG3* cloned into pETDUET-1, C-terminal 10His epitope, ApR | This study |
| pET22b | Expression vector, PBR322 origin, PT7, PelB signal sequence, ApR | Invitrogen |
| pSBC81 | *tle3* cloned downstream the PelB signal sequence in pET22b, C-terminal 10His epitope, ApR | This study |
| pVT3 | *tle4* cloned into pETDUET-1, C-terminal 10His epitope, ApR | This study |
| pBB41 | *PA0262* (△3) cloned into pRSF-DUET, C-terminal Strep epitope, KanR | This study |
| pBB42 | *PA0262* (△2-3) cloned into pRSF-DUET, C-terminal Strep epitope, KanR | This study |
| pBB43 | *PA0262* (1-2-3) cloned into PCDF-DUET, C-terminal 10His epitope, | This study |
| pBB44 | *PA0262* (1-2-3) cloned downstream the PelB signal sequence in pET22b, C-terminal 10His epitope, ApR | This study |
| pBB45 | substitution of the codon coding His935 in Ala of PA0262 in pBB44, C-terminal 10His epitope, ApR | This study |
| pBB46 | substitution of the codon coding Glu936 in Ala of PA0262 in pBB44, C-terminal 10His epitope, ApR | This study |

**Table 2:** Oligonucleotides used in this study.

| Plasmids | Oligonucleotide names and sequences |
|---|---|
| pBB01 | BBO09 : GCAGTGGAACGCCACTGCAGGAACGATAGGGTTCGG |
| | BBO08 : CCGGGGATCCTCTAGATTAGATTGTCCCCCC |
| pBB02 | BBO09 : GCAGTGGAACGCCACTGCAGGAACGATAGGGTTCGG |
| | BBO69 : CCGGGGATCCTCTAGATTAAGCAATGCCGCGCCA |
| pBB03 | BBO70 : GCAGTGGAACGCCACTGCAGGGGGCTCCCGGGAAGG |
| | BBO08 : CCGGGGATCCTCTAGATTAGATTGTCCCCCC |
| pBB04 | BBO12 : GCAGTGGAACGCCACTGCAGGGTTTCCAAACTGCCA |
| | BBO11 : CCGGGGATCCTCTAGATCATGGCTTCTCTCC |
| pBB05 | BBO15 : GCAGTGGAACGCCACTGCAGGGCCGAGTCGCCGCTA |
| | BBO14 : CCGGGGATCCTCTAGACTATAAATCGTCCTG |
| pBB06 | BBO15 : GCAGTGGAACGCCACTGCAGGGCCGAGTCGCCGCTA |
| | BBO67 : CCGGGGATCCTCTAGATTATCGGGGTCGATCCG |
| pBB07 | BBO68 : GCAGTGGAACGCCACTGCAGGCTGGGGCTGGCGCAA |
| | BBO14 : CCGGGGATCCTCTAGACTATAAATCGTCCTG |
| pBB08 | BBO19 : GCAGTGGAACGCCACTGCAGGGCCAACGACGAGTCC |
| | BBO17 : CCGGGGATCCTCTAGATCAAACTGCGGAGGCGGG |
| pBB09 | BBO19 : GCAGTGGAACGCCACTGCAGGGCCAACGACGAGTCC |
| | BBO18 : CCGGGGATCCTCTAGATCAGTATCCCGTTGG |
| pBB10 | BBO07 : GCGCACGCGGCGGGCTGCAGGGAACGATAGGGTTCGG |
| | BBO08 : CCGGGGATCCTCTAGATTAGATTGTCCCCCC |
| pBB11 | BBO10 :GCGCACGCGGCGGGCTGCAGGGGTTTCCAAACTGCCA |
| | BBO11 : CCGGGGATCCTCTAGATCATGGCTTCTCTCC |
| pBB12 | BBO12 : GCGCACGCGGCGGGCTGCAGGGGCCGAGTCGCCGCTA |
| | BBO14 : CCGGGGATCCTCTAGACTATAAATCGTCCTG |
| pBB13 | BBO16 : GCGCACGCGGCGGGCTGCAGGGGCCAACGACGAGTCC |
| | BBO17 : CCGGGGATCCTCTAGATCAAACTGCGGAGGCGGG |
| pBB14 | BBO16 : GCGCACGCGGCGGGCTGCAGGGGCCAACGACGAGTCC |
| | BBO18 : CCGGGGATCCTCTAGATCAGTATCCCGTTGG |
| pBB15 | BBO24 : CCAAGCTTGCATGCCTGCAGATGAACGATAGGGTT |
| | BBO25 : CGGTACCCGGGGATCCTCGATTGTCCCCCCAAA |
| pBB16 | BBO26 : CCAAGCTTGCATGCCTGCAGATGGTTTCCAAACTGCCA |
| | BBO27 : CGGTACCCGGGGATCCTCTGGCTTCTCTCCTTG |
| pBB17 | BBO28 : CCAAGCTTGCATGCCTGCAGATGGCCGAGTCGCCG |
| | BBO29 : CGGTACCCGGGGATCCTCTAAATCGTCCTGCCA |
| pBB18 | BBO39 : CCAAGCTTGCATGCCTGCAGATGGCCAACGACGAG |
| | BBO40 : CGGTACCCGGGGATCCTCAACTGCGGAGGCGG |
| pBB19 | BBO39 : CCAAGCTTGCATGCCTGCAGATGGCCAACGACGAG |
| | BBO41 : CGGTACCCGGGGATCCTCGTATCCCGTTGGGAA |
| pBB20 | BBO24 : CCAAGCTTGCATGCCTGCAGATGAACGATAGGGTT |
| | BBO25 : CGGTACCCGGGGATCCTCGATTGTCCCCCCAAA |
| pBB21 | BBO26 : CCAAGCTTGCATGCCTGCAGATGGTTTCCAAACTGCCA |
| | BBO27 : CGGTACCCGGGGATCCTCTGGCTTCTCTCCTTG |
| pBB22 | BBO28 : CCAAGCTTGCATGCCTGCAGATGGCCGAGTCGCCG |
| | BBO29 : CGGTACCCGGGGATCCTCTAAATCGTCCTGCCA |
| pBB23 | BBO39 : CCAAGCTTGCATGCCTGCAGATGGCCAACGACGAG |
| | BBO40 : CGGTACCCGGGGATCCTCAACTGCGGAGGCGG |
| pBB24 | BBO39 : CCAAGCTTGCATGCCTGCAGATGGCCAACGACGAG |
| | BBO41 : CGGTACCCGGGGATCCTCGTATCCCGTTGGGAA |
| pBB25 | BBO36 : AAGGAGATATACATATGATGCGTCAAAGGGAC |
| | BBO37 : TTTTTCGAACTGCGGGTGGCTCCAAgcgcTGTATCCCGTTGG |
| | BBO34bis : GCGTGGCCGGCCGATATCTCATTTTTCGAACTGCGGGTGGCTCCAAGCGCT |
| pBB26 | BBO05 : AAGGAGATATACATATGGTTTCCAAACTGCCA |
| | BBO34 : TTTTTCGAACTGCGGGTGGCTCCAAgcgcTTGGCTTCTCTCCTTG |
| | BBO34bis : GCGTGGCCGGCCGATATCTCATTTTTCGAACTGCGGGTGGCTCCAAGCGCT |
| pBB27 | BBO06 : TTTTTCGAACTGCGGGTGGCTCCAAgcgcTTGGCTTCTCTCCTTG |
| | BBO35 : TTTTTCGAACTGCGGGTGGCTCCAAgcgcTTAAATCGTCCTGCCA |
| | BBO34bis : GCGTGGCCGGCCGATATCTCATTTTTCGAACTGCGGGTGGCTCCAAGCGCT |
| pBB28 | BBO80 : TTTTTCGAACTGCGGGTGGCTCCAAgcgcTTAAATCGTCCTGCCA |
| | BBO72 : TCAATGTTAATGATGATGGTGATGGTGATGGTGATGATGTAAATCGTCCTGCCAG |
| | BBO71bis : CCGCAAGCTTGTCGATCAATGTTAATGATGATGGTGATGGTGATGGTGATGATG |

| | |
|---|---|
| pBB29 | VTO26 : CAGGTCGACGGATCCCCGGG GGTTGGAGCCGGGAAAGACGT |
|  | VTO27 : GCATTAGATTGTCCCCCCGTTCATGGCTTCTCT |
| | VTO28 : AGAGAAGCCATGAACGGGGGGACAATCTAATGC |
| | VTO29 : TATGCATCCGCGGGCCCGGGGAACAACGCCTCCAGTACTCCTTCAGG |
| pBB30 | BBO22 : CAGGTCGACGGATCCCCGGGCCGGCATGGTTCTGGTCACC |
| | BBO23 : CGCGGGACACCATCGACCTACATTAGATTGTCCCCCCAAA |
| | BBO24 : TTTGGGGGGACAATCTAATGTAGGTCGATGGTGTCCCG |
| | VTO35 : TATGCATCCGCGGGCCCGGGAACGGCGCTCTCCCAGGACC |
| pBB31 | VTO32 : CAGGTCGACGGATCCCCGGGCGCCAAGGTCGATATGAAACACACGA |
| | VTO36 : GCATTAGATTGTCCCCCCATTGGTCAGTATCC |
| | VTO37 : TACTGACCAATGGGGGGGACAATCTAATGCGCT |
| | VTO29 : TATGCATCCGCGGGCCCGGGGAACAACGCCTCCAGTACTCCTTCAGG |
| pBB32 | BBO55 : CAGGTCGACGGATCCCCGGGGAAATGCAAGACCTTGCTCA |
|  | BBO56 : GGACGATTTAAGGGTTAGGGATAGGCTTACCTGGCTTCTCT |
| | BBO57 : CCTATCCCTAACCCTCTCCTCGGTCTCGATTCTACGTGAAAGGAGAGAA |
| | BBO58 : TATGCATCCGCGGGCCCGGGAGCGAGATGAAACCGAACAG |
| pBB33 | BBO52 : CAGGTCGACGGATCCCCGGGGAAGCTCTCACCGCACAACT |
| | BBO53 : ATCGAGACCGAGGAGAGGGTTAGGGATAGGCTTACCTAAATCGTCC |
| | BBO54 : CCTATCCCTAACCCTCTCCTCGGTCTCGATTCTACGTGAGTCGATGGTG |
| | VTO35 : TATGCATCCGCGGGCCCGGGAACGGCGCTCTCCCAGGACC |
| pBB34 | BBO73 : CAGGTCGACGGATCCCCGGGTGGGCACTTTCAACGACAAG |
|  | BBO74 : ATGGTGATGGTGATGATGCCCTGCCACCGGCGAGCGCC |
| | BBO75 : CATCACCATCACCATTGAGTTGACAGGCACACCGGTCG |
| | BBO76 : TATGCATCCGCGGGCCCGGGGTTCCAGCAGCAGTTCGTAG |
| pBB35 | BBO30 : CAGGTCGACGGATCCCCGGGCCGGCATGGTTCTGGTCACC |
| | SBO157 : TGAGAGCATCAAGCGGATTAGATTGTCCCCCCAA |
| | SBO158 : TTGGGGGGACAATCTAATCCGCTTGATGCTCTCA |
| | SBO163 : TATGCATCCGCGGGCCCGGG TGAACAACGCCTCCAGTACT |
| pBB36 | BBO30 : CAGGTCGACGGATCCCCGGGCCGGCATGGTTCTGGTCACC |
| | SBO159 : ACTGGCAAAGCAGCGGATCGAGGATAGCGGCGA |
| | SBO160 : TCGCCGCTATCCTCGATCCGCTGCTTTGCCAGT |
| | SBO163 : TATGCATCCGCGGGCCCGGG TGAACAACGCCTCCAGTACT |
| pBB37 | BBO30 : CAGGTCGACGGATCCCCGGGCCGGCATGGTTCTGGTCACC |
| | SBO161 : ATAAGCATCAGCGGAGATGGTGGCCTCCTTGGC |
| | SBO162 : GCCAAGGAGGCCACCATCTCCGCTGATGCTTAT |
| | SBO163 : TATGCATCCGCGGGCCCGGG TGAACAACGCCTCCAGTACT |
| pBB38 | BBO42 : GCTTGATATCGAATTCGAGGTCGGACTATCC |
| | BBO46 : TCGGCGTTGGGATACGTTCCTTGTCGTTG |
| | BBO47 : AACGTATCCCAACGCCGAGGAGCGGCAAG |
| | BBO48 : CGGGCTGCAGGAATTAGATTGTCCCCCC |
| pBB39 | BBO42 : GCTTGATATCGAATTCGAGGTCGGACTATCC |
| | BBO43 : GTTTTTCAGGGATACGTTCCTTGTCGTTG |
| | BBO44 : AACGTATCCCTGAAAAACTTCCCAACGGG |
| | BBO45 : CGGGCTGCAGGAATTCTCATGGCTTCTCTCC |
| pBB40 | BBO42 : GCTTGATATCGAATTCGAGGTCGGACTATCC |
|  | BBO49 : ACCGCCTGCGGATACGTTCCTTGTCGTTG |
| | BBO50 : AACGTATCCGCAGGCGGTGCGCTTTGGGG |
| | BBO51 : CGGGCTGCAGGAACTATAAATCGTCCTG |
| pBB41 | BBO36 : AAGGAGATATACATATGATGCGTCAAAGGGAC |
| | BBO107 : GCGTGGCCGGCCGATATCTCATTTTTCGAACTGCGGGTGGCTCCAAGCGCTAACTGCGGAGGCGGG |
| pBB42 | BBO36 : AAGGAGATATACATATGATGCGTCAAAGGGAC |
| | BBO103 : TTTTTCGAACTGCGGGTGGCTCCAAGCGCTGGCCGCGCCGACATG |
| | BBO37bis : GCGTGGCCGGCCGATATCTCATTTTTCGAACTGCGGGTGGCTCCAAgcgcT |
| pBB43 | BBO93 : AAGGAGATATACATATGATGGAGCACCAGGGCGTG |
| | BBO94 : GCGTGGCCGGCCGATATCTCAATGTTAATGATGATGGTGATGGTGATGGTGATGATG |
| pBB44 | BBO86 : CCGGCGATGGCCATGGGGGAGCACCAGGGCGTG |
| | BBO87 : ATGTTAATGATGATGGTGATGGTGATGGTGATGATGGTATCCCGTTGGGAA |
| | BBO87bis : GGTGGTGGTGCTCGAGTCAATGTTAATGATGATGGTGATGGTGATGGTGATGATG |
| pBB45 | BBO88 : CAGGGTCTGTTCATTGCCGAAATGACCCATGTC |
| | BBO89 : GACATGGGTCATTTCGGCAATGAACAGACCCTG |
| pBB46 | BBO95 : GGTCTGTTCATTCACGCCATGACCCATGTCTGG |
| | BBO96 : CCAGACATGGGTCATGGCGTGAATGAACAGACC |
| pVT1 | VTO01 : AGGAGATATACCATGGATGAACGATAGGGTT |
| | VTO02 : CCGCAAGCTTGTCGACTCAATGTTAATGATGATGGTGATGGTGATGGTGATGATGGATTGTCCCCCCAAA |
| pVT3 | VTO10 : AGGAGATATACCATGGATGAACGATAGGGTT |
| | VTO11 : CCGCAAGCTTGTCGACTCAATGTTAATGATGATGGTGATGGTGATGGTGATGATGTGTCCCTCCTTT |
| pVT8 | VTO06 : AAGGAGATATACATATGAAATGCAAGACCTTG |
|  | VTO07 : GCGTGGCCGGCCGATATCTCACGTAGAATCGAGACCGAGGAGAGGGTTAGGGATAGGCTTACCTGGCTTCTCTCCTTG |
| pVT9 | VTO08 : AAGGAGATATACATATGCGCTTGATGCTCTCA |
| | VTO09 : GCGTGGCCGGCCGATATCTCACGTAGAATCGAGACCGAGGAGAGGGTTAGGGATAGGCTTACCTAAATCGTCCTGCCA |
| pSBC81 | VTO51 : CCGGCGATGGCCATGGGGAACGATAGGGTTCGG |
| | VTO52 : GGTGGTGGTGCTCGAGGATTGTCCCCCCAAA |
| pSBC107 | CSO31 : CTCGATTCTACGTGAGATATCTTTAAGAAGGAGATATACATAT |
| | CSO32 : GCGTGGCCGGCCGATATCTCAGTGGTGGTGGTGGTGGTG |

### Supplementary Materials and Methods

Fractionation of *E. coli* cells into spheroplasts (cytoplasm and membranes) and periplasmic fractions were done as described previously [8].

**Supplementary figure 1**

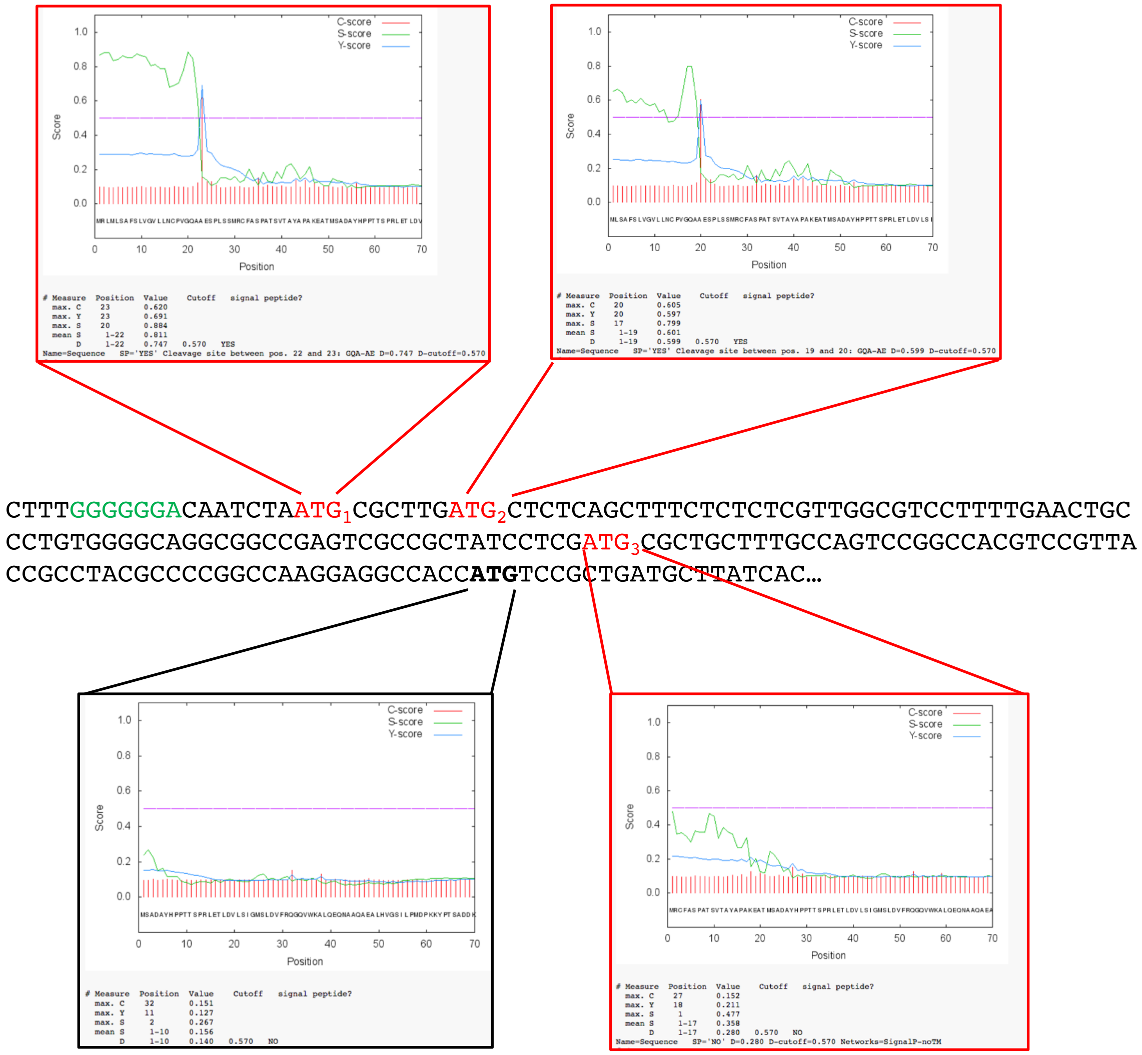

**Fig. S1 : *In silico* analysis of the nucleic and protein sequence of Tla3 (PA0259).** For the analysis of the PA0259 sequence, three translation initiation codons (in red) upstream of the annotated ATG on the genome (in bold) and in phase with the PA0259 sequence were taken into consideration. The green sequence represents a putative Shine-Dalgarno sequence for the $ATG_1$ initiation codon. SignalP 4.1 program was used to locate signal sequence at the N-terminus of the fours corresponding proteins. The program predicts the residue where the cleavage site (C-score) takes place, the most probable position of the signal sequence (S-score) and combines these two approaches (Y- and D-scores).

**Supplementary Figure 2**

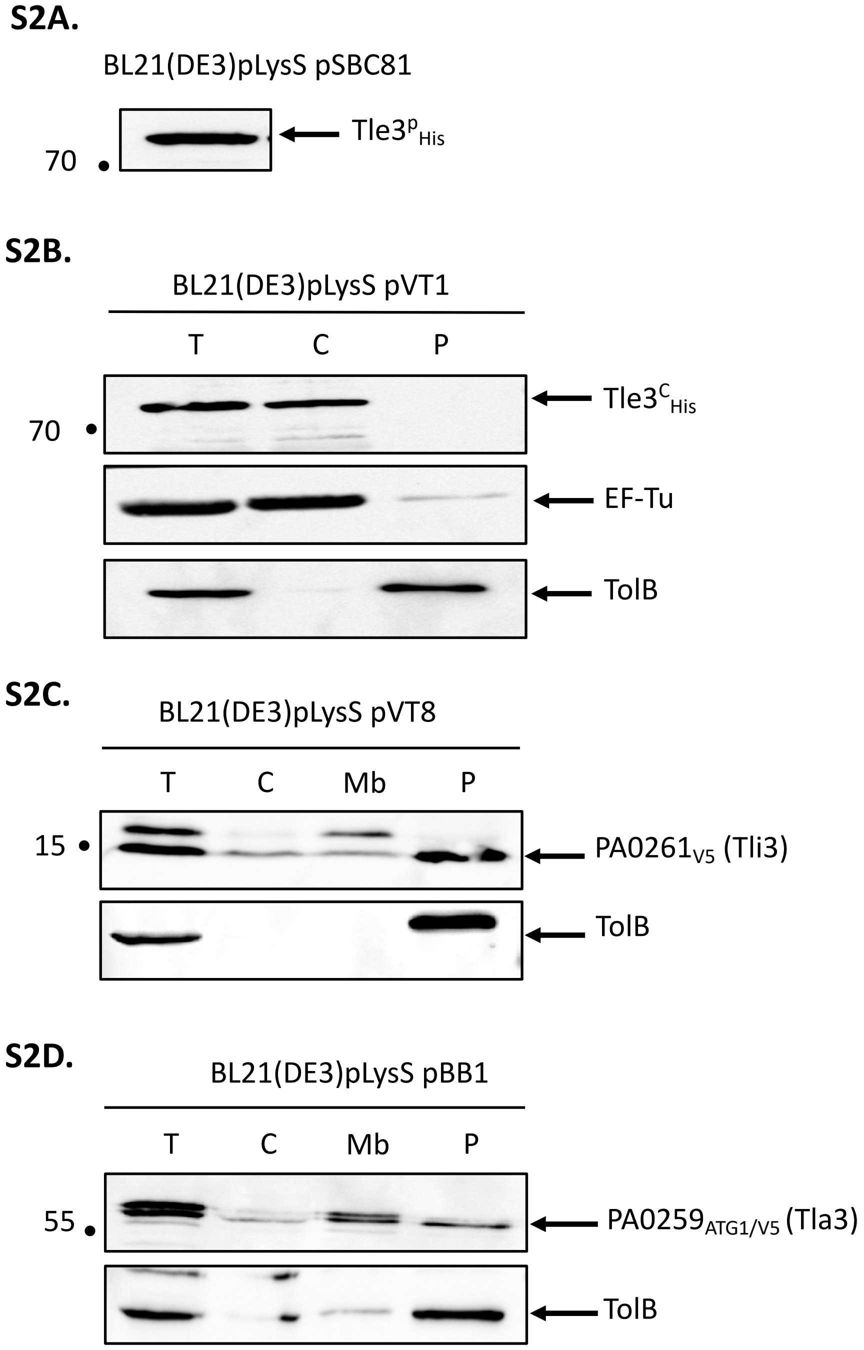

**Fig. S2 : Production of Tle3$^{p}$ (S2A) and localization of Tle3$^{c}$, Til3 and Tla3$_{ATG1}$ in *E. coli* BL21(DE3)pLysS (S2B-D).** Except for Tle3$^{P}$ whose production promotes cell lysis, bacteria were subjected to fractionation and immunoblotting using antibodies directed against the His Tag, V5 tag, EF-Tu and TolB. EF-Tu and TolB were used as cytoplasmic and periplasmic controls respectively. T: whole cell, C: cytoplasm, Mb: total membrane, P: periplasm. The position of the proteins and the molecular mass markers (in kDa) are indicated.

**Supplementary figure 3**

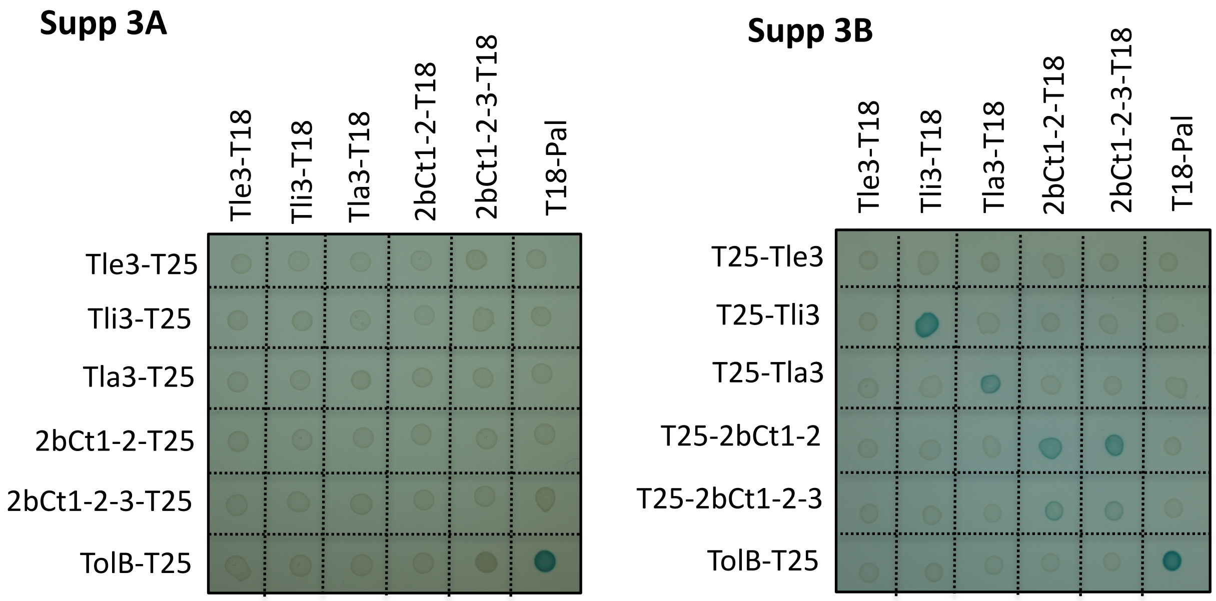

**Fig. S3: Bacterial two-hybrid assay (A and B)**. BTH101 reporter cells producing the indicated proteins or domains fused to the T18 or T25 domain of the *Bordetella* adenylate cyclase were spotted on X-gal indicator plates. The blue color of the colony reflects the interaction between the two proteins. TolB and Pal are two proteins known to interact but unrelated to the T6SS. The experiment was performed in triplicate and a representative result is shown.

**Supplementary figure 4**

**S4A.**

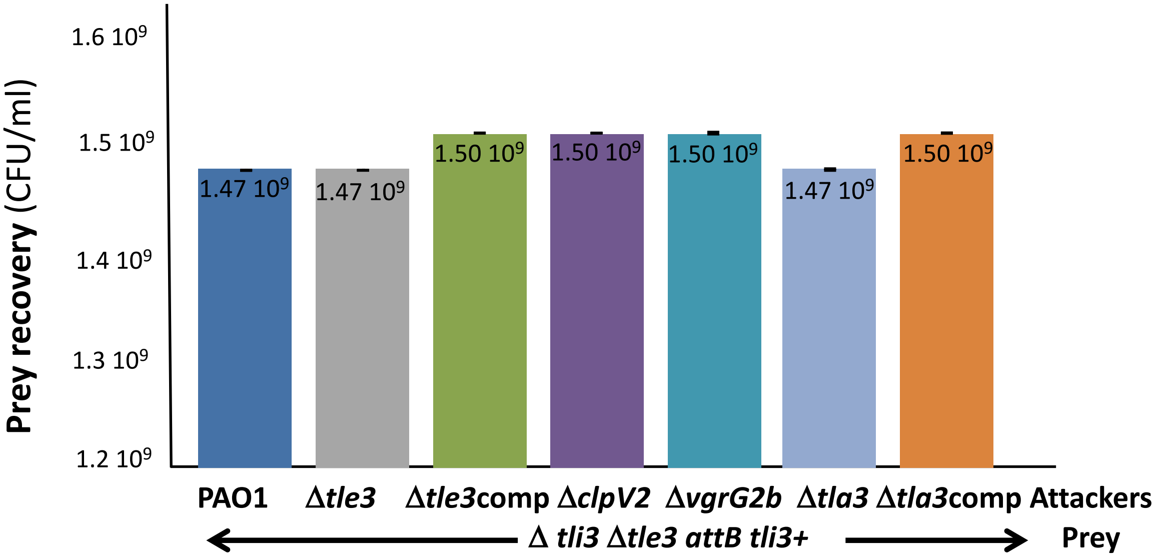

**S4B.**

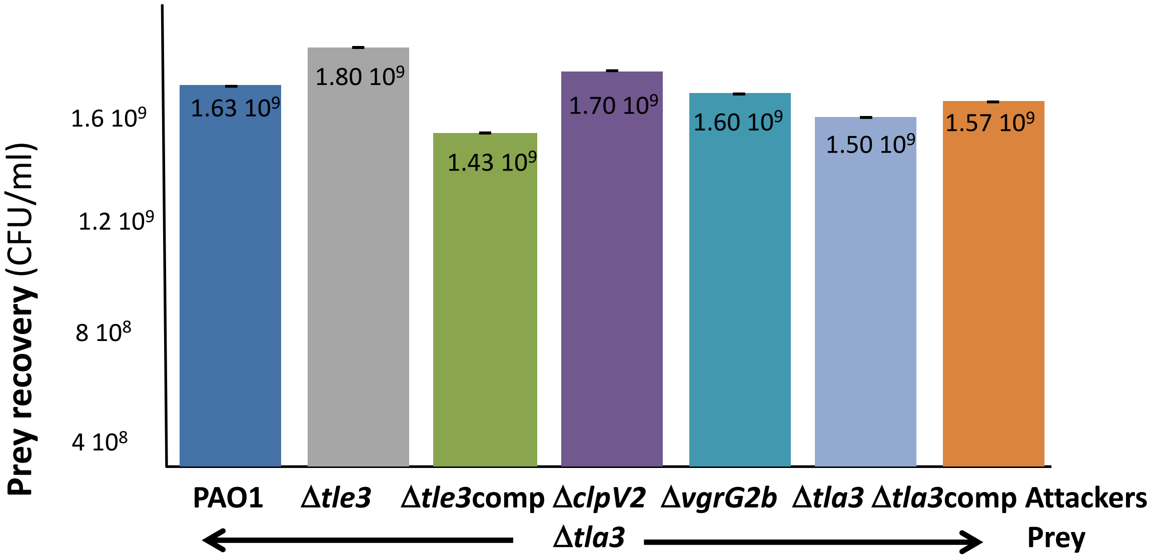

**Fig. S4: *P. aeruginosa* growth competition.** The *P. aeruginosa* prey strain Δ*tli3*Δ*tle3 attB tli3+* **(S4A)** or Δ*tla3* **(S4B)** was incubated with various *P. aeruginosa* attacker strains as indicated in the figure for 24h at 37°C. The number of recovered prey bacteria is indicated in CFU/ml. "comp" stand for *cis* complementation of the corresponding mutation with a wild-type copy inserted at the *attB* site on *P. aeruginosa* chromosome. Error bars represent ± SEM (n = 3)

Supplementary figure 5

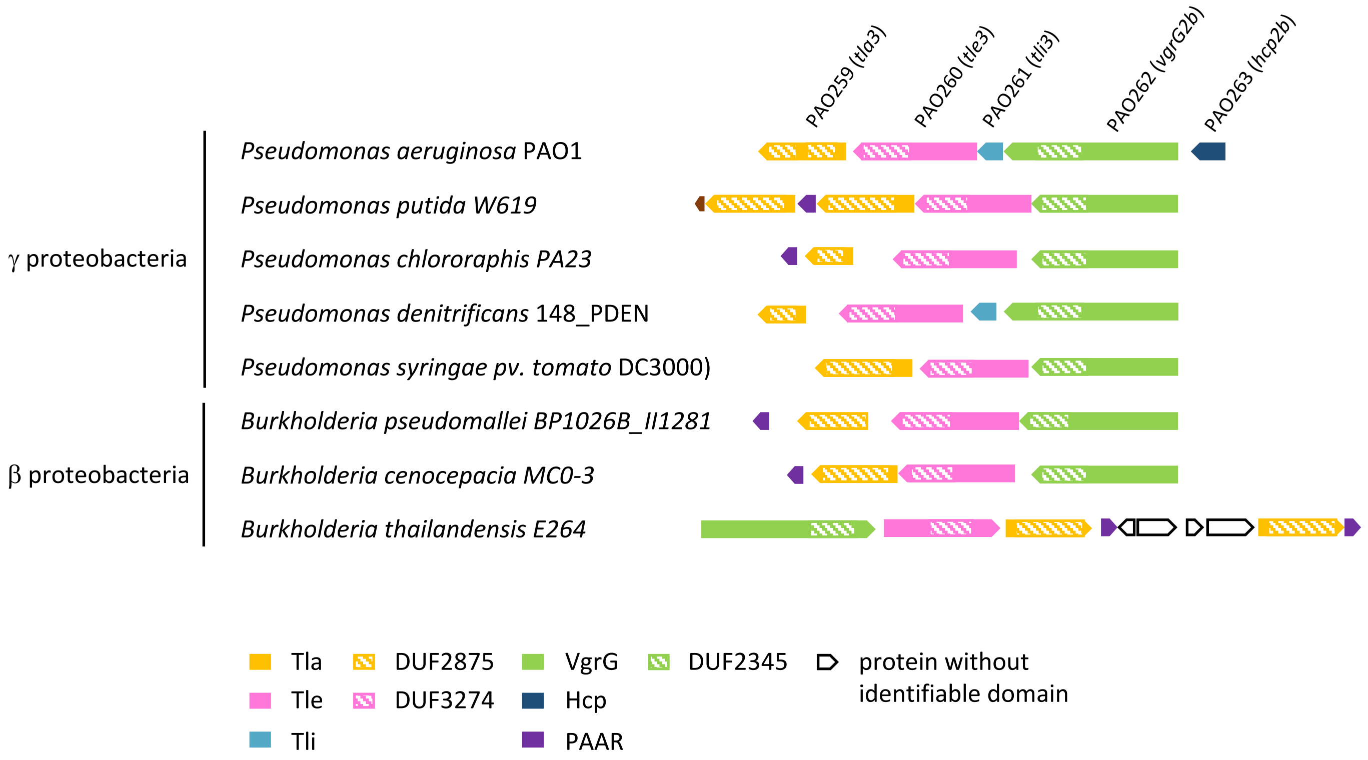

Fig. S5: **Adaptors of Tla3 family DUF2875 are restricted to beta and gamma proteobacteria.**
Tla3 homologues were found in bacteria belonging to γ and β proteobacterial classes. Genetic organization presented at right of each bacterium was deduced from manual inspection of respective bacterial genome and color-coded by the presence of a conserved protein domain as predicted by an NCBI-CD search algorithm. The names of *P. aeruginosa* genes are indicated and the color-coded protein domains are listed and drawn to scale.
